# A remarkable transformation catalyzed by a domain-of-unknown-function 692 during the biosynthesis of a new RiPP natural product

**DOI:** 10.1101/2023.02.06.527370

**Authors:** Richard S. Ayikpoe, Lingyang Zhu, Jeff Y. Chen, Chi P. Ting, Wilfred A. van der Donk

## Abstract

The domain of unknown function 692 (DUF692) is an emerging family of posttranslational modification enzymes involved in the biosynthesis of ribosomally-synthesized and posttranslationally modified peptide (RiPP) natural products. Members of this family are multinuclear iron-containing enzymes and only two members have been functionally characterized to date: MbnB and TglH. Here, we used bioinformatics to select another member of the DUF692 family, ChrH, that is ubiquitously encoded in the genomes of the *Chryseobacterium* genus along with a partner protein ChrI. We structurally characterized the ChrH reaction product and show that the enzyme catalyzes an unprecedented chemical transformation that results in the formation of a macrocycle, an imidazolidinedione heterocycle, two thioaminals, and a thiomethylation. Based on isotopic labeling studies, we propose a mechanism for the four-electron oxidation and methylation of the substrate peptide. This work identifies the first SAM-dependent DUF692 enzyme, further expanding the repertoire of remarkable reactions catalyzed by these enzymes.

## Introduction

Recent advances in bioinformatic tools and an explosion in the publicly available genomic data have led to the identification of many new peptide-based natural products.^1–6^ Often, these compounds possess antimicrobial, antiviral, antifungal, herbicidal or cytotoxic properties.^7–11^ Ribosomally synthesized and posttranslationally modified peptides (RiPPs) have garnered significant attention over the last decade owing to their remarkable structural diversity and biological activities.^12,13^ RiPPs are produced from genetically encoded precursor peptides that are extensively modified by posttranslational modification enzymes encoded in the same biosynthetic gene clusters (BGCs). Many of the post-translational modification reactions are unique chemical transformations resulting in macrocycles and heterocycles, structures that are privileged in bioactive scaffolds. Furthermore, exciting recent developments in biocatalysis have increasingly shown the value of enzymatic transformations in preparation of compounds that are useful to human society, with many of the enzymes coming from biosynthetic pathways to natural products.^14–17^ Therefore, discovery of new chemical reactions catalyzed by enzymes and assignment of function to poorly characterized enzyme families is an important goal.

A common biosynthetic feature of most RiPPs is the ribosomal production of a precursor peptide consisting of a leader and a core sequence. The leader peptide facilitates recognition and processing by the posttranslational modification (PTM) enzymes,^18^ while modifications are imparted to the core peptide. The modified peptide undergoes proteolytic removal of the leader peptide, and the matured peptide is in most cases exported from the cell to exert its biological function. More than 40 families of RiPPs have been classified based on the chemical transformations made to the precursor peptide.^12,13^ One emerging and underexplored group of RiPPs are generated by multinuclear iron-dependent enzymes belonging to the DUF692 family. Members of this enzyme family are structurally related to the triose-phosphate isomerase family,^19^ and only two members of the DUF692 enzyme family have been functionally characterized thus far, MbnB^20–24^ and TglH. ^23,24^

MbnB forms a heterodimeric complex with MbnC to catalyze a central step in the biosynthesis of methanobactin, a copper-chelating peptidic compound produced by methanotrophic bacteria under copper limiting conditions. During the maturation of methanobactin, MbnBC catalyzes the four-electron oxidation of Cys residues in its precursor peptide, MbnA, to an oxazolone and an adjacent thioamide group (Figure 1A).^20–22^ The second characterized member, TglH is involved in the maturation of 3-thiaglutamate, an amino acid-derived natural product biosynthesized at the C-terminus of a carrier peptide.^23^ TglH, forms a complex with a second protein TglI to catalyze the β-carbon excision of a C-terminal Cys residue to generate a 2-mercaptoglycine residue (Figure 1A).^23–25^ Although they belong to the same enzyme family, TglH and MbnB catalyze completely different but equally remarkable chemical transformations on their substrate peptides. This precedent motivated us to investigate additional members of the DUF692 family. Herein, we used bioinformatics to uncover a new RiPP BGC that encodes a DUF692 homolog. This BGC is encoded in the genomes of several members of the *Chryseobacterium* genus. We therefore propose the name chryseobactin for the natural product of the pathway. Structural investigations of the product of the DUF692 enzyme revealed an unprecedented chemical transformation distinct from those catalyzed by the two previously characterized members.

**Figure 1.**
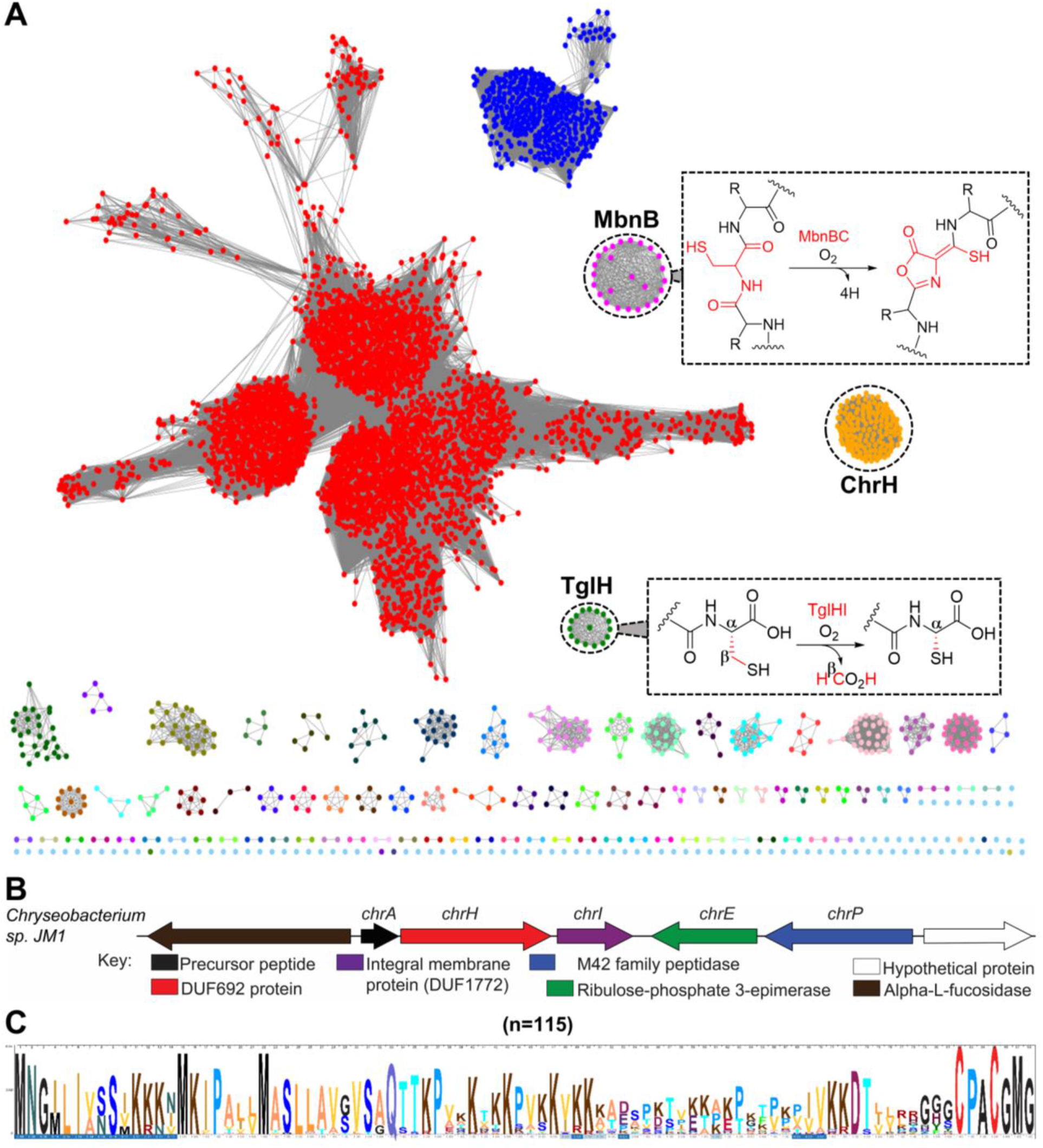
Bioinformatic analysis of the DUF692 family. (A) Sequence similarity network of DUF692 enzymes using an E-value of 50. The orthologs of MbnB are colored green and the orthologs of TglH in magenta. The group of BGCs investigated in this work are colored orange. (B) Representative gene organization of the *Chryseobacterium* biosynthetic gene clusters. (C) Multiple sequence alignment of the precursor peptides showing conservation of a terminal CPACGMG motif.

## Results and Discussion

### RiPP biosynthetic pathways that encode uncharacterized DUF692 enzymes

To identify additional members of the DUF692 family, we generated a sequence similarity network (SSN) using the Enzyme Function Initiative Enzyme Similarity Tool (EFI-EST).^26^ A total of 13,136 UniRef90 sequences of the PF05114 family were used in the initial SSN analysis. For the SSN shown in Figure 1, an E-value of 10^-50^ and a sequence alignment score threshold of 70 were used resulting in more than 100 groups. These groups were colored based on the gene context of each cluster, and analyzed using the EFI’s genome neighborhood network (GNN) tool.^27^ The analysis identified clusters that are associated with the two characterized members TglH and MbnB (Figure 1A). In addition, the analyses also revealed additional clustered family members including a group of enzymes in the *Chryseobacterium* genus. We selected this group of BGCs for further investigation because of the gene organization and the sequences of the putative precursor peptides (Figure 1B and 1C), which suggested products distinct from methanobactin and 3-thiaglutamate.

This group of BGCs encodes proteins annotated as a precursor peptide (ChrA), a DUF692 enzyme (ChrH), a DUF1772-containing integral membrane protein (ChrI), a ribulose phosphate 3-epimerase (ChrE), and an M42 peptidase (ChrP). Multiple sequence alignments of the precursor peptides from 115 BGCs revealed a highly conserved CPACGMG motif at the C-terminus (Figure 1C). While our bioinformatic discovery was based on an enzyme SSN followed by GNN analysis, similar BGCs were also recently identified using a co-occurrence based approach.^28^

### ChrH requires ChrI to modify the CPACGMG motif of ChrA

The two previously characterized members of the DUF692 family both modify cysteine residues on their substrates. We therefore anticipated that the conserved Cys residues in ChrA might be modified by ChrH. To probe this hypothesis, we employed a heterologous coexpression approach to characterize the activity of ChrH. The precursor peptide ChrA was expressed in *Escherichia coli* with an N-terminal His6-tag. Purification of ChrA by immobilized metal affinity chromatography (IMAC) and subsequent analysis by matrix-assisted laser desorption/ionization time-of-flight mass spectrometry (MALDI-TOF MS) gave the expected mass for the unmodified linear peptide (Figure 2B). When ChrA was coexpressed with ChrH in lysogeny broth (LB) supplemented with 50 μM iron(II) citrate, no change in the mass of the purified peptide was observed.

**Figure 2.**
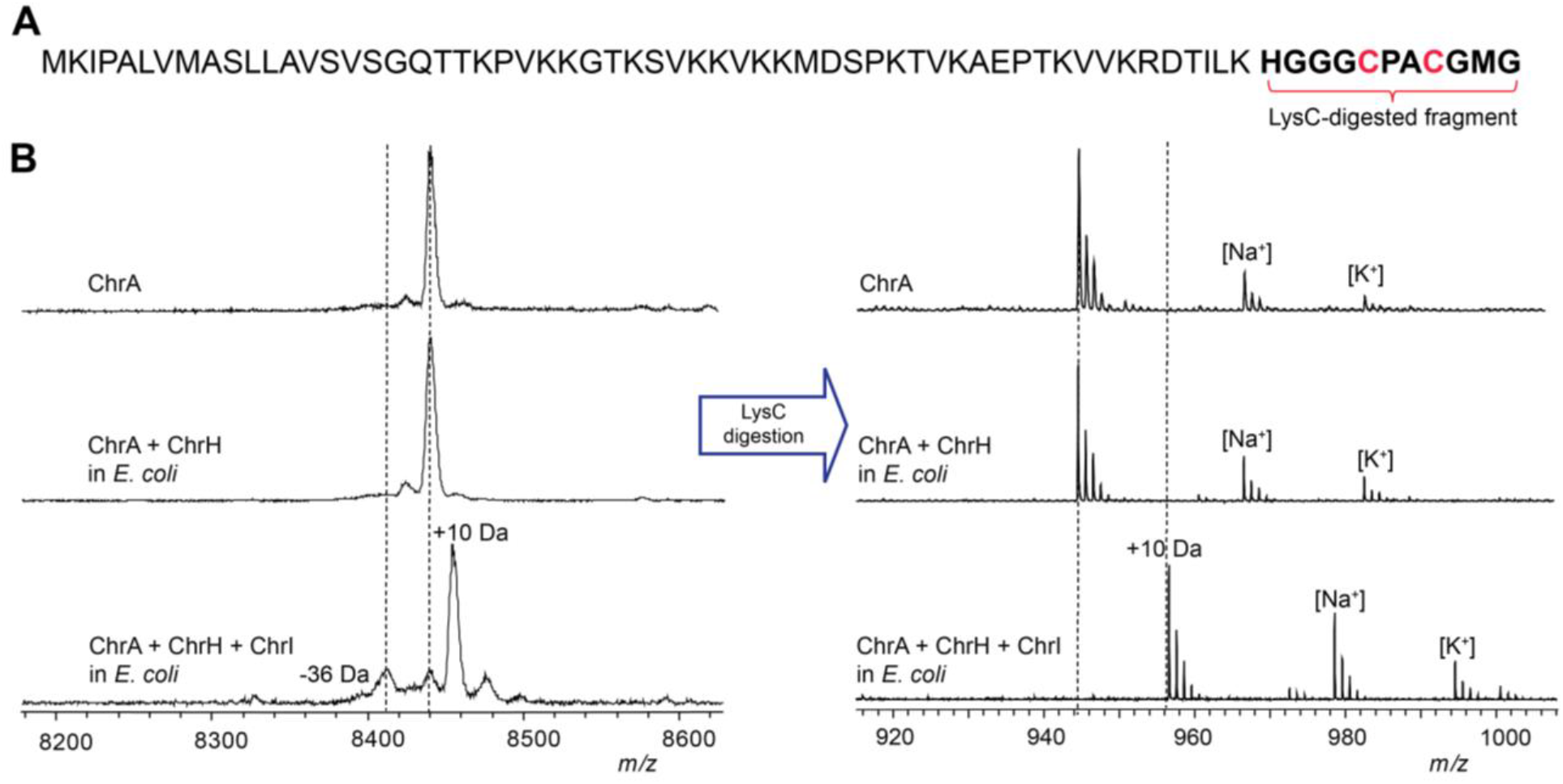
Modification of ChrA by ChrHI in *E. coli.* (A) Sequence of ChrA showing the C-terminal fragment after digestion with endoproteinase LysC. (B) MALDI-TOF MS spectra of the unmodified and modified full-length ChrA (left) and the corresponding LysC-digested peptides (right).

Both MbnB and TglH require a second protein for activity.^20,23^ MbnB forms a heterodimer with MbnC,^21,22^ and TglH forms a predicted heterodimer with TglI.^24^ Both MbnC and TglI recognize the leader peptides of their cognate substrates by formation of an antiparallel β-sheet,^21^ with TglI having the typical fold of a RiPP Recognition Element (RRE).^29^ The integral membrane protein ChrI is not homologous in sequence to MbnC or TglI and was not bioinformatically predicted to contain an RRE,^30^ nor do any of the other enzymes in the *chr* BGC. ChrI does contain a DUF1772 that has been speculated to play a role in protein-protein interactions.^31^ We therefore next included ChrI in the expression system. Following coexpression of all three proteins, a new peak consistent with a 10 Da increase in mass of ChrA was observed by MALDI-TOF MS (Figure 2B). Digestion of isolated modified and unmodified peptide with endoproteinase LysC suggested the modification is localized to the C-terminus of ChrA (Figure 2B). The 10 Da mass increase was confirmed by high-resolution mass spectrometric (HRMS) analysis (Figure S1 and S2). We designate this LysC-digested peptide ChrA*.

To determine if the conserved Cys residues were modified, we first derivatized ChrA and ChrA* with *N*-ethylmaleimide (NEM) to probe for free thiols. Analysis of the derivatized peptides by MALDI-TOF MS showed that ChrHI-processed ChrA did not react with NEM (Figure S3), suggesting that the thiols have been modified. To corroborate this conclusion, the two conserved Cys residues in ChrA, Cys63 and Cys66 were mutated to Ser residues. Neither single nor double mutants resulted in formation of the product that had increased by 10 Da, suggesting that both Cys residues are modified by ChrH (Figure S4). The ChrA-C6S variant resulted in partial production of a product with a decrease in mass of 36 Da that will be further discussed below. Taken together, these results suggest that ChrH, like TglH and MbnB, requires a complex with a second protein to install posttranslational modifications on ChrA and that this transformation involves the thiols of the two conserved Cys residues of ChrA.

### Structure elucidation of ChrA*

To gain more detailed structural information regarding the reaction product of ChrHI, both unmodified and modified ChrA were produced on a larger scale and subjected to detailed one-dimensional and two-dimensional (1D and 2D) NMR analysis after endoproteinase LysC digestion to reduce the size to an 11mer peptide. Analysis of TOCSY data for LysC-digested ChrA identified all 11 amino acids including nine amide protons (Figure S5 and Table S1), whereas only seven spin systems were identified for ChrA* associated with just seven amide protons (Figures 3A, S6 and Table S2). Significantly, the amide protons of Gly9 and Met10 (Gly67 and Met68 of full-length peptide) were missing in ChrA*, suggesting that these two amides were modified (Figures 3A and S6). Subsequent ^1^H-^13^C-HSQC analysis of substrate and product also revealed a new cross peak with chemical shifts at 1.96 (^1^H) and 10 ppm (^13^C) (Figure S6). Interestingly, this new peak integrated in the ^1^H spectrum to three protons and appeared as a singlet suggesting the possible introduction of a methyl group. The ^1^H and ^13^C chemical shifts of this methyl group suggest attachment to either carbon or a heteroatom like sulfur that is not highly electronegative.

**Figure 3.**
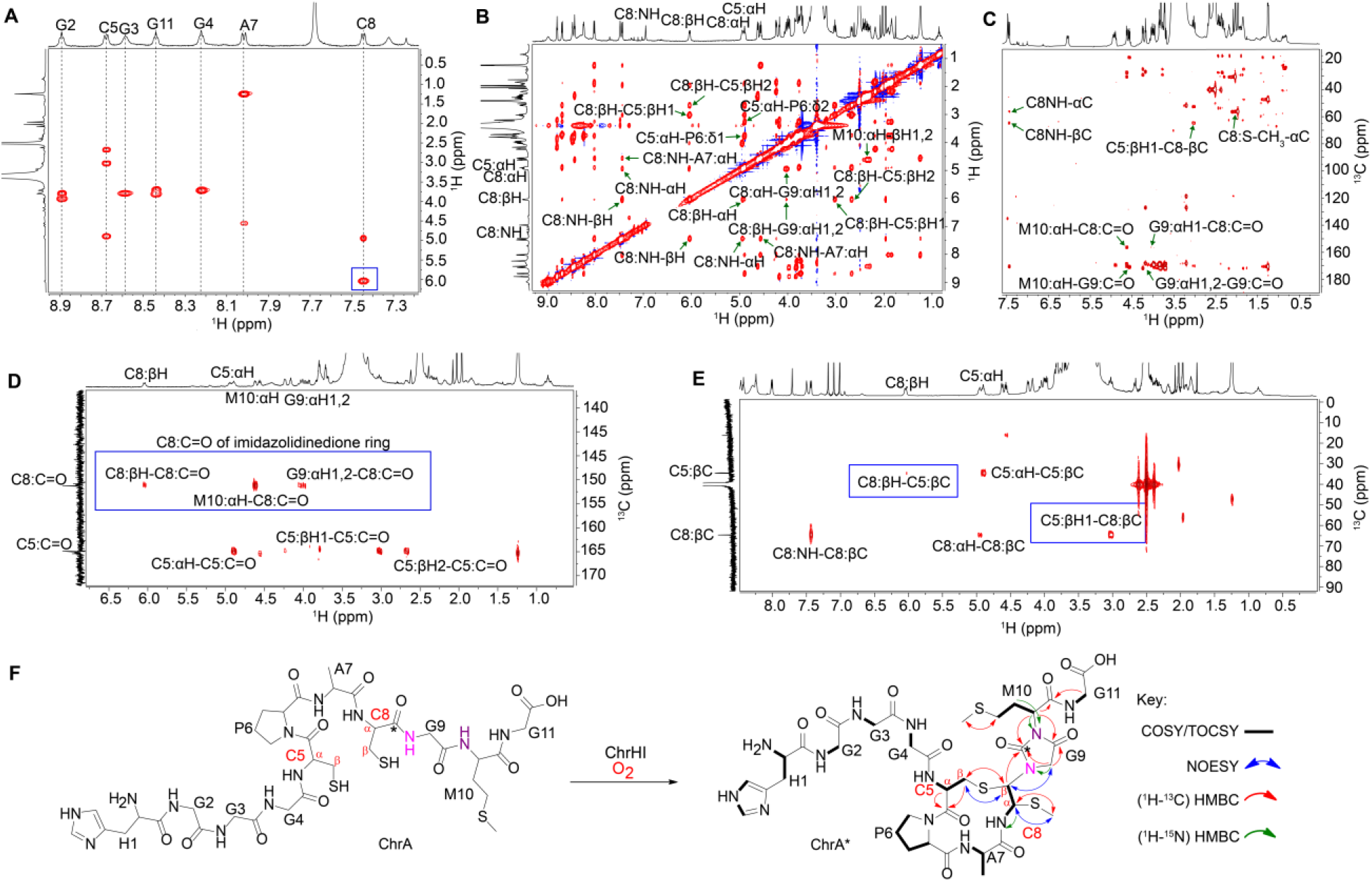
NMR analysis of ChrA*. (A) TOCSY spectrum showing the loss of amide NH signals of Gly9 and Met10 and a change in the chemical shift of βH of Cys8 (blue box). (B) NOESY spectrum focusing on important NOE correlations in the macrocycle and heterocycle of ChrA*. (C) HMBC spectrum highlighting the through-bond correlations of Cys8-Cα-S-CH_3_, and the correlations that establish the macrocycle and the heterocycle of ChrA*. (D) HMBC spectrum of 1-^13^C-Cys labeled ChrA* showing the connectivities of the Cys8 carbonyl carbon with resonances derived from Cys8, Gly9, and Met10 (blue box) that establish the imidazolidinedione. (F) HMBC spectrum of 3-^13^C-Cys labeled ChrA* highlighting the connectivity between the βH of Cys5 and the βC of Cys8 (blue box) as well as an important connectivity between the macrocycle and heterocycle. (F) Proposed structure of ChrA* showing all important correlations identified from the NMR spectral analysis.

In addition, the HSQC data identified the disappearance of the CH2 group at the β position of the former Cys8. Instead, a new CH cross peak was observed in ChrA* at 6.04 ppm, a significant change in the chemical shift compared to the original β-protons of ChrA (3.36 and 2.96 ppm; Figure S6 and Tables S1 and S2). As we will show, this new signal arises from one of the β-protons of the former Cys and we will refer to it as such in the remainder of the discussion. The chemical shift of the αH of the former Cys8 also changed from 4.44 ppm in ChrA to 4.94 ppm in ChrA* (Figures S6 and Tables S1 and S2). The assignments of the new signals originating from the α and β protons of the former Cys8 in ChrA* were supported by the ^1^H-^1^H dqCOSY spectrum, which showed cross-peaks between the proton at 4.94 ppm and the NH of the former Cys8, and between the two protons at 4.94 and 6.04 ppm (Figure S6).

To probe the chemical environment of the new signals further, ^1^H-^13^C-HMBC and ^1^H-^1^H NOESY experiments were carried out on ChrA*. The NOESY spectra showed direct correlations between the new methyl group and the αH of the former Cys8. In addition, a second NOE correlation was observed from the βH of Cys5 to the signal at 6.04 ppm associated with the βH of the former Cys8 (Figures 3B and S6), as well as a weaker nOe between the βH of Cys5 and the signal at 4.94 ppm associated with the αH of the former Cys8. The HMBC data also showed connectivity (via 3-bond correlations) between the new methyl group and the αC of the former Cys8 and vice versa. Another connectivity between the βC of Cys5 and the ^1^H peak at 6.04 ppm (former βH of Cys8) was observed, suggesting the formation of a macrocycle. The HMBC spectrum also revealed direct connectivities between the carbonyl of the former Cys8 (156.5 ppm) and the αH of Gly9 and αH of Met10, indicating a significant rearrangement in ChrA* (Figure 3C).

These preliminary experiments suggested that Cys8 is significantly altered. To probe the fate of the individual carbon atoms of Cys8, we first prepared ChrA* generated from ChrA that was selectively labeled with 1-^13^C-Cys (label at the carbonyl carbon) for HMBC NMR analysis. As expected, two carbon peaks, one at 170.0 ppm for Cys5 and the other at 156.5 ppm for the former Cys8 were observed in the ^13^C NMR spectrum (Figure S7), corroborating a significant rearrangement of the position of the carbonyl carbon of the former Cys8. The HMBC spectrum also revealed direct correlations between the βH of the former Cys8 to the carbonyl of the former Cys8, the αH of Met10 to the carbonyl carbons of Gly9 as well as the former Cys8, and from the αH of Gly9 to the carbonyl carbons of Gly9 and the former Cys8 (Figures 3D and S7). When ChrA* was labeled with 3-^13^C-Cys (label at the β–carbon), the acquired HMBC data showed through-bond connectivities from the βC of Cys5 to the βH of the former Cys8 and vice versa, suggesting that the sulfur of Cys5 is directly connected to the Cβ of the former Cys8 (Figures 3E and S8). Collectively, all collected NMR data are consistent with the structure shown in Figure 3F. The stereochemistry at the two new stereogenic centers of the former Cys8 is currently not known. However, the ^3^*J*H-H coupling constant between the protons at 4.94 and 6.04 ppm is 11.6 Hz, suggesting that the relative stereochemistry involves a *trans* arrangement.

To corroborate the findings from the NMR data, ChrA* was next analyzed by high-resolution electrospray ionization tandem mass spectrometry (HR-ESI MS/MS). The HR-ESI MS/MS spectrum localized the modification to the C-terminal CGMG peptide as no fragments corresponding to b-ions could be observed beyond this point, whereas all amino acids in the N-terminal segment were unmodified (Figure 4). The observation of fragment ions indicated as “y4” and “b7” (since the structure is no longer an α-amino acid peptide these fragments are not true y and b ions) is explained by cleavage of the thioaminal linkages in the proposed structure (Table S3). In addition, various internal fragment ions consistent with the proposed structure of ChrA* were observed, especially fragmentation in the two thioether bridges as well as the amide bonds of the imidazolidinedione. Taken together, the HR-ESI MS/MS and the NMR data suggest that ChrHI catalyzes an unprecedented multistep chemical transformation on ChrA: macrocyclization to form a crosslink between the thiol of Cys5 and the βC of the former Cys8, and migration of the carbonyl carbon of the former Cys8 to become inserted between the amide nitrogens of Gly9 and Met10 to form an imidazolidinedione. Furthermore, the reaction results in the installation of a thiomethyl group at the αC of the former Cys8. Thus, the net +10 Da transformation from ChrA observed by MS involves a −4 Da change resulting from two oxidative ring formations in which the amide protons of Gly9 and Met10 are removed as well as the thiol proton of Cys5 and one of the β-protons of the former Cys8, and a + 14 Da change from the methylation event.

**Figure 4.**
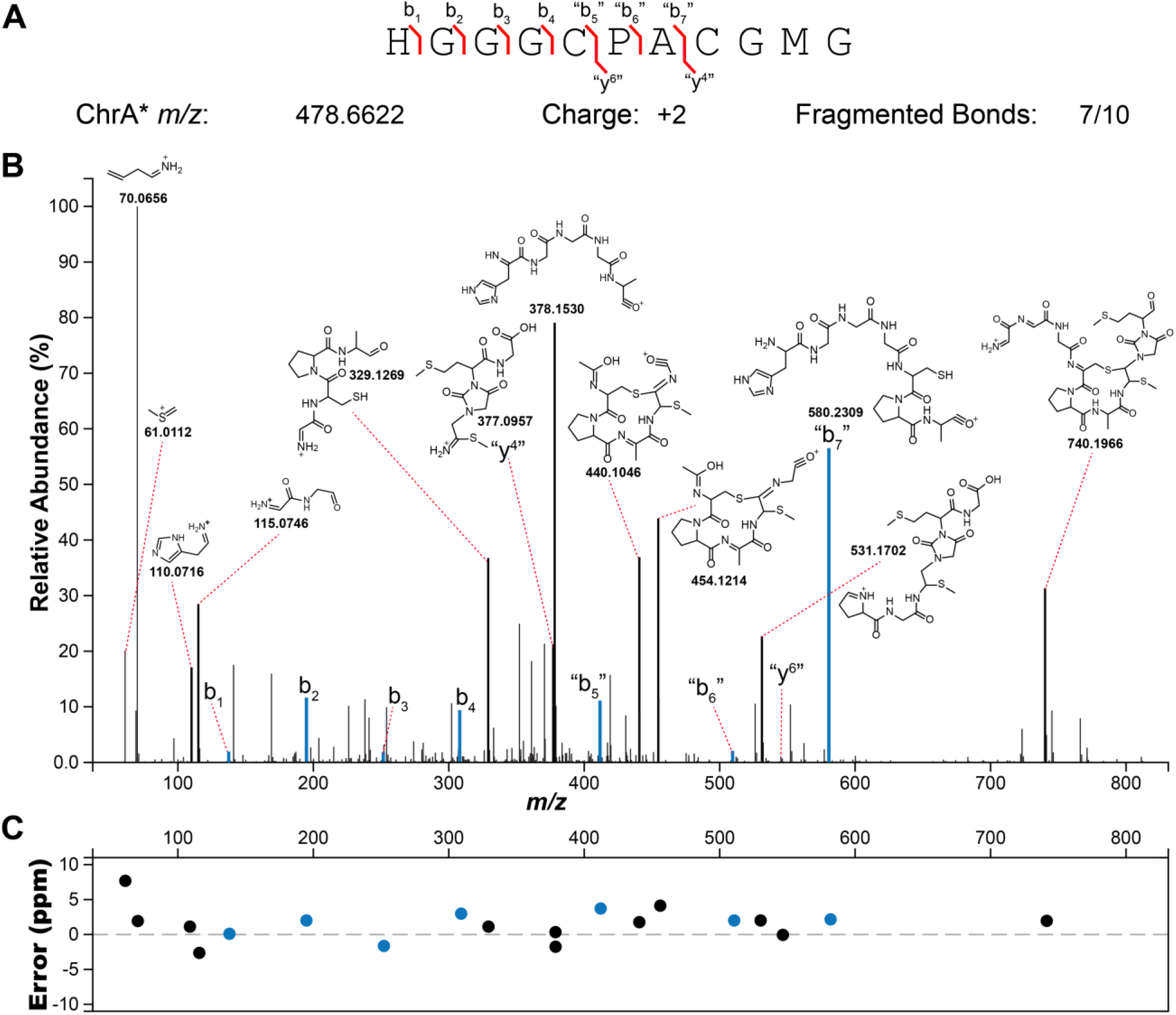
HR-ESI tandem mass spectrometry analysis of ChrA* digested with LysC. (A) Sequence of ChrA* and observed fragment ions. (B) Tandem mass spectrum showing ions and their corresponding fragments. (C) A graph of the ppm errors for each identified ion.^32^

### In vitro reconstitution of ChrHI activity and identification of S-adenosylmethionine as the methyl donor

The proposed structure of ChrA* generated in *E. coli* contains a new methyl group that was not present in the ribosomally generated precursor peptide. This finding raised the question regarding the source of the methyl group. To determine its origin, isotope feeding experiments were carried out. ChrA was coexpressed with ChrHI in *E. coli* in M9 minimal media supplemented with ^13^CD_3_-Met.^33^ Isolation of the modified peptide and subsequent MALDI-TOF MS analysis of the full-length and LysC-digested peptide revealed the incorporation of two methyl groups from ^13^CD_3_-Met into ChrA*, one in Met10 and one in the new thiomethyl group (Figures 5A and S9). When selenomethionine was substituted for ^13^CD_3_-methionine, the isolated product contained four selenium atoms in the full length modified ChrA consistent with the four Met residues in ChrA and only one selenium atom in the ChrA* as evidenced by the isotope distribution pattern (Figures 5A and S10).^34^ These results suggest that only the methyl group (and not the thiomethyl group) of methionine is incorporated into ChrA* and suggests that either *S*-adenosylmethionine (SAM) or possibly a methionine residue of the modifying enzyme ChrH, could be the methyl donor.

**Figure 5.**
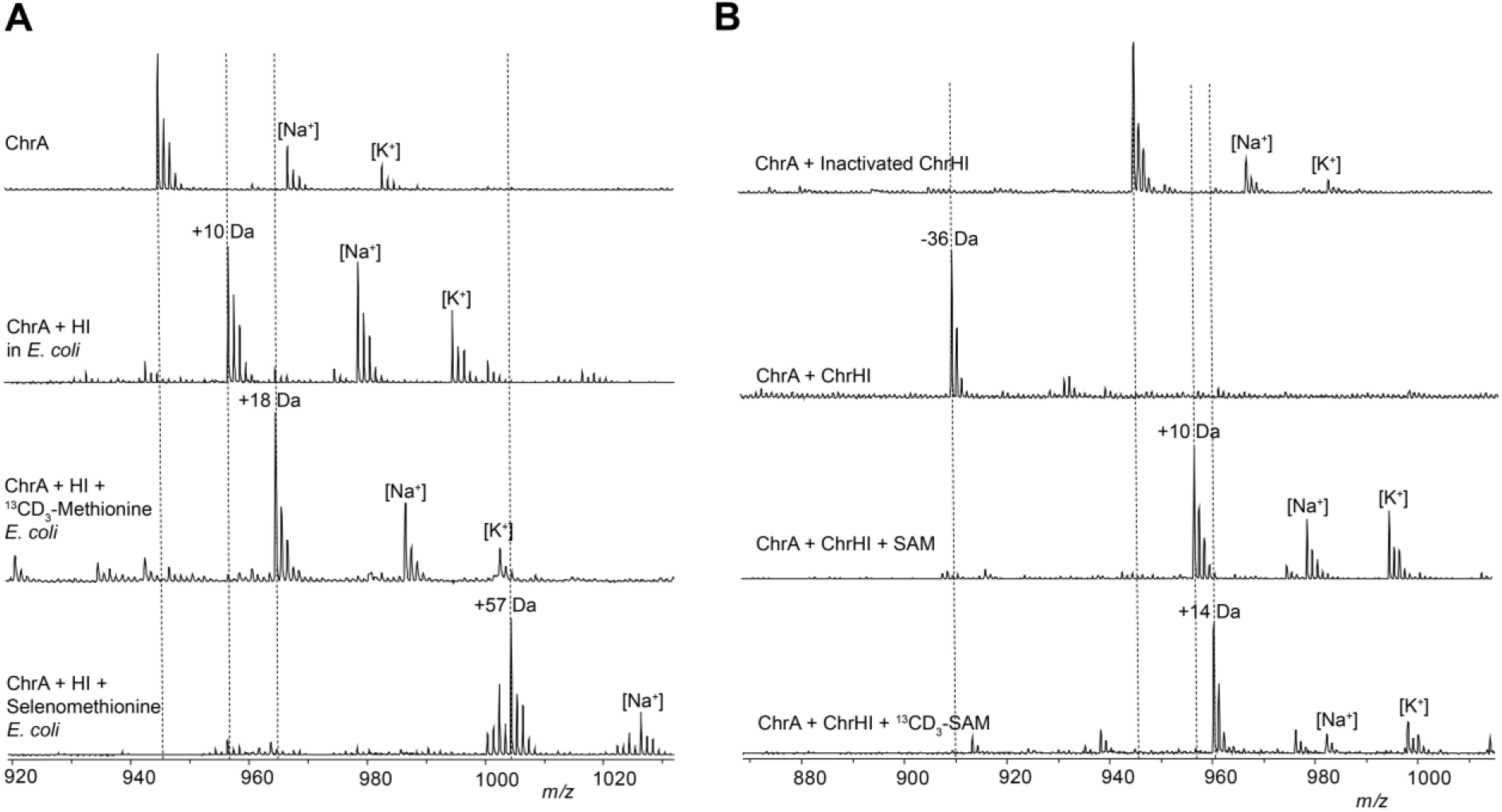
Determination of the methyl source and *in vitro* reconstitution of ChrHI activity. (A) MALDI-TOF MS data of ChrA* produced by heterologous coexpression of ChrA with ChrHI in M9 media supplemented with either ^13^CD_3_-Met or selenomethionine and treatment of the purified peptide with endoproteinase LysC. (B) MALDI-TOF MS of LysC-digested *in vitro* reaction products of ChrHI in the presence and absence of SAM. The dashed lines indicate the expected *m/z* of modified and unmodified peptides.

SAM is the most common methyl donor in biology.^35^We next aimed to reconstitute the activity of ChrHI *in vitro* to probe for the possibility of SAM as the methyl source. ChrH was heterologously expressed in and aerobically purified from *E. coli* RosettapLysS cells as an N-terminally His6-tagged protein (Figure S11A). The purified protein exhibited a purple color. The amount of iron was quantified using the ferene assay,^36^ and the as-isolated enzyme contained 1.9 equivalents of iron per monomer of ChrH. Since the coexpression studies revealed the requirement of ChrI for ChrH activity, the predicted integral membrane protein ChrI was also heterologously expressed in *E. coli* as an N-terminally hexahistidine-tagged protein and partially purified in the presence of detergent (0.01% n-dodecyl-β-D-maltopyranoside) (Figure S11A). Reactions were carried out aerobically by incubating ChrA with ChrH, ChrI and DTT in the presence and absence of SAM. When the reaction without SAM was analyzed by MALDI-TOF MS, the +10 Da product was not observed, but instead, an ion indicating the loss of −36 Da from ChrA was observed (Figures 5B and S12). This new species was also observed in small amounts when ChrA was heterologously coexpressed with ChrHI in *E. coli* (Figure 2B and S3). The difference in mass between the −36 Da species and +10 Da product is 46 Da, consistent with a thiomethyl substituent. Further attempts to isolate the −36 species for detailed structural characterization proved unsuccessful due to rapid degradation under purification conditions. We next analyzed the *in vitro* reaction in the presence of SAM resulting in the formation of the + 10 Da peptide as main product together with the −36 Da species (Figure 3B and S12). These results suggest that SAM is indeed the source of the methyl group and shows that *in vitro* and in *E. coli* the same product is formed strongly suggesting that the observed transformation is the native function of ChrHI. We next synthesized ^13^CD_3_-SAM enzymatically,^37–39^ and used the isotopically labeled product in the *in vitro* reaction with ChrA and ChrHI. Analysis of the full-length and LysC-digested reaction products revealed the incorporation of one ^13^CD_3_-methyl group from ^13^CD_3_-SAM into ChrA*, but no label incorporation in the – 36 Da product. Taken together, these results suggest that SAM is the methyl donor in the posttranslational modification of ChrA by ChrHI thus representing the first member of the DUF692 family to use SAM and iron as cofactors.

### Possible mechanism of ChrHI catalysis

With the structure of ChrA* as well as the source of the methyl group established, a proposed mechanism for the ChrHI-catalyzed transformation of ChrA is shown in Figure 6. DUF692 enzymes have been shown to contain two or three iron ions in their active sites (Protein Database accession numbers 3BWW, 7DZ9, 7FC0, 7TCR, 7TCX, 7TCU, and 7TCW).^20–23^ In the structure of MbnB bound to its partner MbnC and its substrate MbnA, the side chain sulfur atom of a Cys in the substrate is liganded to one of the iron atoms.^21^ The protein ligands to bind two or three iron ions are conserved in ChrH and present in the active site of an AlphaFold model of the enzyme (Figure S13). Akin to TglH^23^ and MbnB,^21,22^ we suggest that the active form of ChrH requires at least two iron ions, one of which is in the Fe(II) form that is used for catalysis and a second (or third) ion that is in the Fe(III) oxidation state that is important for substrate binding and positioning. Such use of iron ions in different oxidation states for different roles is an emerging feature of multinuclear mixed-valent non-heme iron enzymes.^40–44^ It is possible that both Cys thiols ligate to different iron ions in the active site, facilitated by the Pro-Ala sequence that separates the two Cys residues that is an inducer of β-turn formation. Because we currently do not have any structural data on the coordination of the substrate to the metal ions, the mechanism in Figure 6 shows just one of the Cys side chains ligated to Fe as in the MbnABC co-crystal structure.^21^

The Fe^II^ ion in the active site of ChrH will react with molecular oxygen to form a superoxo-Fe^III^ intermediate **I**,^22,45,46^ which initiates the reaction by abstracting a hydrogen atom from the β-carbon of Cys8 as in the proposed mechanisms for TglH and MbnB.^20–23^ The highly reducing thioketyl radical formed during this abstraction is expected to transfer an electron to one of the Fe(III) ions to generate a thioaldehyde and Fe(II). As drawn in Figure 6, this is a different iron than the iron that activates O2 but it could be the same iron (e.g. some of the mechanisms shown in Figure S14). Subsequent attack of the amide nitrogen of Gly9 onto the thioaldehyde forms a β-lactam **II**, possibly facilitated by the iron ion functioning as a Lewis acid. Formation of a similar β-lactam was also proposed for MbnBC catalysis as well as other mononuclear non-heme iron enzymes such as isopenicillin N synthase.^20,22,47–49^ At this point, the reaction of ChrHI diverges from previous enzymes in that a second nucleophilic attack of the amide nitrogen of the downstream Met10 onto the carbonyl of the β-lactam would result in the formation of a bicyclic intermediate **III**. For MbnBC, the β-lactam is attacked by the oxygen atom of the upstream amide,^21^ possibly with the intermediacy of an enzyme bound intermediate.^22^

The transfer of an electron from Fe(II) in intermediate **II** to the ferric hydroperoxo intermediate results in generation of a ferryl (Fe^IV^-oxo) species. The exact timing of the formation of this highly reactive intermediate is interesting as it could be favorable to delay its formation until after (or concomitant with) the formation of the bicycle in intermediate **III**. One possible means to delay its formation could be by having the reducing equivalent residue on an iron atom that is not bound to the peroxide as shown in Figure 6. Once formed, the ferryl could initiate a proton-coupled electron transfer from the adjacent O-H bond to generate an oxygen radical **IV**. A well-precedented β-scission reaction would result in the opening of the four-membered ring to form a resonance stabilized carbon-based radical **V**.^50^ Although the context and enzyme classes are quite different, β-scission by oxidation of an alcohol by a ferryl is also proposed as a key step in other non-heme iron dependent enzymes and experimental and computational support has been provided.^41,51,52^ We propose that this intermediate radical **V** could form the product via formation of an episulfide followed by nucleophilic ring opening by Cys5 and methylation of the thiol (pathway A). Alternatively, oxidation of radical **V** to a cation **VI** (pathway B) followed by *S-*methylation of the thioaminal, migration to the Cβ of the former Cys and addition of the thiol of Cys5 to the Cα of the former Cys8 would complete the formation of ChrA*. We note that many of these steps could occur in a different order, that several steps are expected to require acid-base catalysis, and that other mechanisms can be drawn to arrive at the final product (see Figure S14).

The mechanism in Figure 6 (and alternatives in Figure S14) may also explain the formation of the −36 Da species detected in our assays. Unlike MbnB and TglH, ChrH also uses SAM as a cofactor. *In vitro* reconstitution of ChrHI revealed that in the absence of SAM, only a −36 Da species accumulates as evidenced by MALDI-TOF MS (Figure 5 and S12). This compound cannot be generated by elimination of methanethiol from ChrA* as that would lead to a product that would be 34 Da lighter than ChrA. We propose that in the absence of SAM, radical **V** does not form the episulfide (pathway A) or the cation **VI** (pathway B). Instead, the radical **V** could be reduced leading to the product shown in pathway C (Figure 6). Regardless of the exact mechanism, methylation appears critical to arrive at the structure of ChrA*.

The mechanisms in Figures 6 and S14 can potentially also explain the outcome with the ChrA variants C63S and C66S. Since the proposed chemistry is initiated at Cys66, replacement with a Ser residue abolishes all activity. In contrast, replacement of Cys63 with Ser can still result in the initial oxidation steps on Cys66. If macrocycle formation involving Cys63 is required for either methylation or thiomethyl migration to the former α-carbon then once again a shunt product can be formed that is 36 Da decreased in mass from the starting peptide (Figure S15)

**Figure 6.**
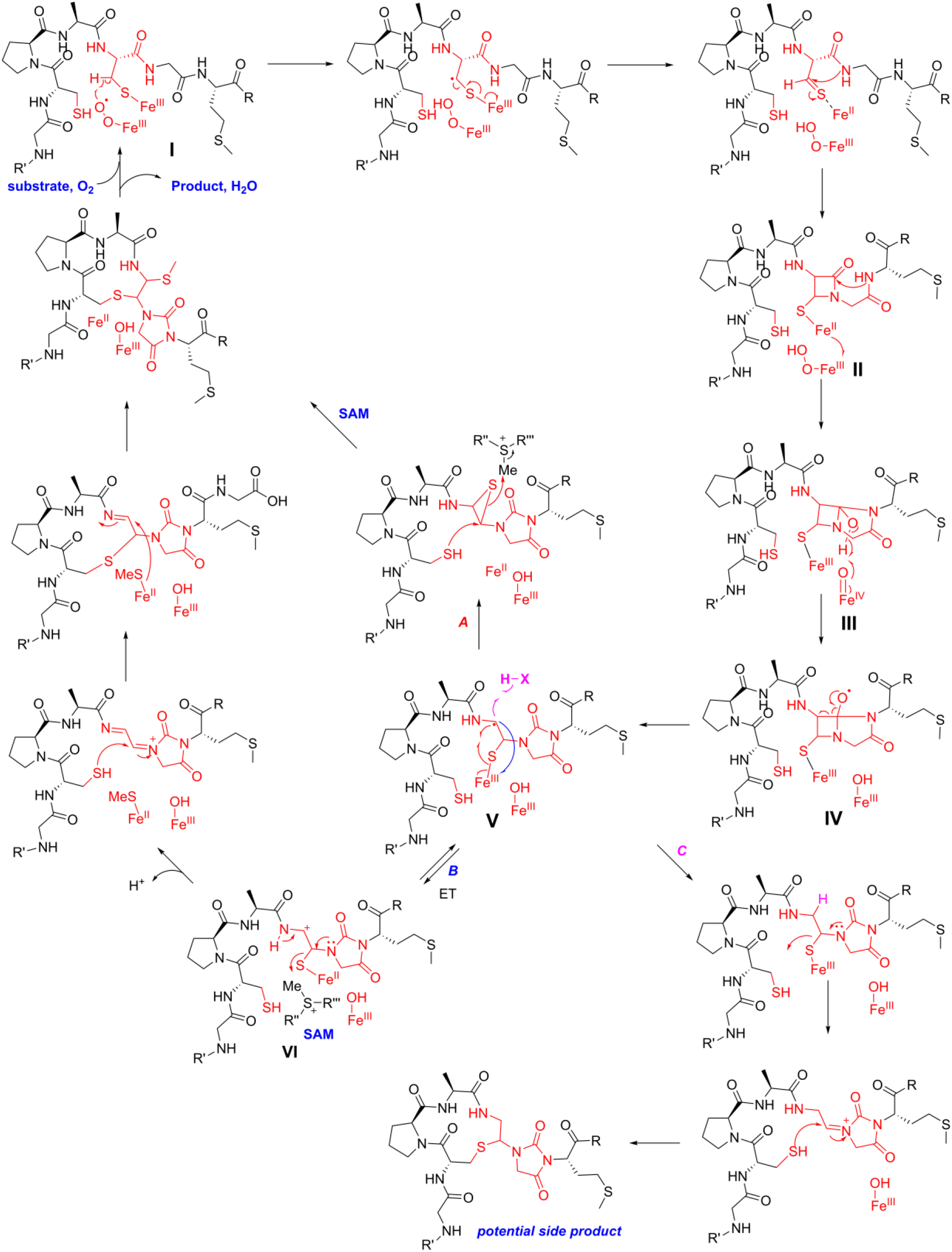
Proposed mechanism for ChrHI catalysis. The order of some steps can be inverted (e.g. the two C-S bond formatting steps) and the timing of methylation is not known and could happen much earlier (e.g. in intermediate **II** or **III**). For some alternative mechanisms, see Figure S14.

## Conclusion

Using bioinformatics, we identified many groups of uncharacterized enzymes that belong to the DUF692 enzyme family. By combining biochemical assays, mass spectrometric and spectroscopic experiments, we demonstrate that one group of such enzymes catalyzes a peptide backbone rearrangement to form an imidazolidinedione heterocycle adding to the repertoire of heterocycles formed in RiPPs. In addition to heterocycle formation, ChrH installs a thioether macrocycle from two conserved Cys residues and methylates a thiohemiaminal using SAM. Future spectroscopic studies and possibly structural information on the observed side products may provide further insights into the mechanism of the remarkable overall process.

The final structure of chryseobactin will require reconstitution of the remaining two enzymes encoded in the BGC, a protease ChrP and a putative epimerase ChrE, which has not yet been achieved to date. Their characterization or that of orthologs would facilitate investigation into the biological function of chryseobactin. Collectively, the data in this study expand the scope of posttranslational modifications in RiPP biosynthesis, demonstrate another unexpected and complex reaction catalyzed by a DUF692 enzyme, and lay the foundation towards understanding the chemistry of additional members of the DUF692 enzyme family.

## Funding

This work was supported by the National Institutes of Health (F32 GM140621 to RSA and R37 GM058822 to WAV).

## Notes

The authors declare no competing financial interest(s).

## Acknowledgement

This work is dedicated to the memory of Prof. Christopher T. Walsh whose pioneering studies on enzyme mechanisms and natural product biosynthesis have been an inspiration to the authors. We thank D. T. Nguyen for his assistance with HR-MS/MS experiments. We also thank Dr. P. Yau and Dr. J. Arrington in the Roy J. Carver Biotechnology Center at the University of Illinois at Urbana-Champaign for assistance with MS/MS data acquisition.

## Notes

### Competing Interest Statement

The authors have declared no competing interest.

