## Supplemental Information for "A remarkable transformation catalyzed by a domain-of-unknown-function 692 during the biosynthesis of a new RiPP natural product"

The data in this study has been deposited in: van der Donk, Wilfred; Ayikpoe, Richard; Zhu, L (2023), "Data associated with DUF692 enzyme from Chryseobacterium", Mendeley Data, V1, doi: 10.17632/dn37cj6z5m.1. The data will be released upon publication.

### Table of Contents

**Materials and methods** ... 2

**Figure S1.** High-resolution electrospray ionization mass spectrometry (HRMS) analysis of LysC-digested ChrA and ChrA* ... 8

**Figure S2.** Characterization of LysC-digested ChrA and ChrA* ... 9

**Figure S3.** Characterization of modified ChrA. ... 10

**Figure S4.** Mutational analysis of ChrA ... 11

**Figure S5.** NMR spectra of LysC-cleaved ChrA peptide in 90% $H_2O$ and 10% $D_2O$ at 25 $^oC$ ... 14

**Figure S6.** NMR spectra of LysC-cleaved ChrA* peptide in DMSO-d6 at 23 $^oC$. ... 21

**Figure S7.** NMR spectra of LysC-cleaved 1-$^{13}C$-Cys-labeled ChrA* peptide in DMSO-d6 at 23 $^oC$ ... 23

**Figure S8.** NMR spectra of LysC-cleaved 3-$^{13}C$-Cys-labeled ChrA* peptide in DMSO-d6 at 23 $^oC$. ... 26

**Figure S9.** Determination of the methyl source for ChrHI catalysis. ... 27

**Figure S10.** MALDI-TOF-MS analysis of selenoMet-ChrA*. ... 28

**Figure S11.** Preparation of ChrH and ChrI. ... 28

**Figure S12.** In vitro reconstitution of ChrHI. ... 29

**Figure S13**. Predicted structure of ChrH using AlphaFold. ... 30

... 32

**Figure S14.** Alternative mechanisms for ChrHI catalysis ... 33

**Figure S15.** Possible explanation for the formation of a -36 Da product from the ChrA-C63S variant ... 34

**Table S1.** Chemical Shifts of ChrA in 90% $H_2O$ and 10% $D_2O$ at 25 $^oC$. ... 35

**Table S2.** Chemical Shifts of ChrA* in DMSO-d6 at 23 $^oC$. ... 35

#### Materials and methods

Primers and enzymes used for molecular biology work were purchased from Integrated DNA Technologies Inc. (Coralville, IA) and New England Biolabs (Ipswich, MA), respectively. Reagents, buffer components and LB were purchased from Fischer Scientific (Hampton, NH). Isopropyl β-D-1-thiogalactopyranoside (IPTG), ampicillin, streptomycin and kanamycin were purchased from Gold Biotechnology (Olivette, Missouri). $^{13}$C- Cysteine and $^{13}CD_3$-methionine were purchased from Cambridge Isotope Laboratories, Inc (Tewksbury, MA). Gel extraction and plasmid isolation were conducted using QIAprep spin columns according to the manufacturer's protocol (Qiagen, DE). DNA sequencing was performed by ACGT Inc. (Wheeling, IL) and Plasmidsaurus (Eugene, OR). *E. coli* NEB 5 alpha chemically competent cells were used for cloning and plasmid propagation, while *E. coli* BL21 (DE3) and RosettapLysS were used for peptide expression and protein expression.

##### *Construction of pACYCDuet_His$_6$chrA*

The gene encoding ChrA was identified in the genome of *Chryseobacterium* sp. JM1 (Uniprot ID: IW22_14840) and the codon-optimized gene for expression in *E. coli* was purchased as a synthetic gene fragment from Twist Bioscience with 5' and 3' extensions containing overlaps for Gibson assembly. The vector pACYCDuet was linearized with BamHI and HindIII restriction enzymes to allow for gene insertion into the multiple cloning site (MCS) 1. The linearized vector was added to the *chrA* gene fragment in a ratio of 1:2 and assembled using NEBuilder HiFi assembly kit following the manufacturers protocol. The ligation mixture was used to transform NEB 5 alpha chemically competent cells and selected on LB agar plate supplemented with 33 µg/mL of chloramphenicol. Single colonies were selected and used to inoculate 5 mL of LB supplemented with appropriate antibiotic and grown at 37 $^{o}$C overnight. Plasmids were isolated via QIAprep Spin Miniprep Kit (Qiagen) and inserts were verified by whole plasmid sequencing (Plasmidsaurus).

##### *Construction of pET28a_His$_6$chrI and pET28a_His$_6$chrH*

Protein sequences for each gene in the *chr* biosynthetic gene cluster were identified from *Chryseobacterium* sp. JMI (Uniprot IDs: IW22_14850 and IW22_14845). The codon-optimized genes were purchased as synthetic gene fragments from Twist Biosciences with 5' and 3' terminal extensions containing necessary overlaps for Gibson assembly. The pET28A(+) vector was digested with NdeI and XhoI restriction enzymes and the digestion mixture was separated on 0.75% agarose gel. Bands corresponding to the linearized vector were excised and purified using the Qiagen gel extraction kit. Gene fragments were combined with linearized vectors in a ratio of 2:1 and ligated using NEBuilder HiFi assembly kit following the manufacturer's protocol. The ligation mixture was used to transform NEB 5 alpha chemically competent cells and selected on LB agar plates containing 50 µg/mL of kanamycin by overnight incubation at 37 $^{o}$C. Single colonies were selected and used to inoculate 5 mL of LB with appropriate antibiotic and cells were grown at 37 °C overnight. Target vectors were isolated from the overnight cultures using the QIAprep Spin Miniprep Kit (Qiagen) and verified by whole plasmid sequencing (Plasmidsaurus).

***Construction of pCDFDuet_chrHI***

The codon-optimized genes encoding *chrH* and *chrI* were amplified from the plasmids pET28a_His6*chrH* and pET28a_His*chrI*, respectively, using NEB Q5 high fidelity polymerase. The PCR product and NcoI/SacI-digested pCDFDuet vector were separated on a 0.75% agarose gel. Bands corresponding to *chrH*, *chrI* and linearized pCDFDuet were excised and extracted with Qiagen gel extraction kit. The *chrH* gene fragment was combined with linearized pCDFDuet vector in a ratio of 2:1 and the plasmid was assembled using the NEBuilder HiFi assembly kit following the manufacturer's protocol. The assembly mixture was used to transform NEB 5 alpha chemically competent cells and transformants were selected on LB agar plates supplemented with 50 µg/mL of streptomycin. Single colonies were used to inoculate LB containing appropriate antibiotics. The plasmids were isolated using QIAprep Spin Miniprep Kit (Qiagen) and the sequence verified by whole plasmid sequencing for the DNA insert. The sequence-verified pCDFDuet_*chrH* plasmid was linearized with MfeI and XhoI restriction enzymes, separated on a 0.75% agarose gel and purified. The linearized pCDFDuet_*chrH* plasmid was combined with *chrI* gene fragment in a ratio of 1:2 and ligated using the NEBuilder HiFi assembly kit. The ligation mixture was used to transform NEB 5 alpha chemically competent cells and transformants were selected on LB agar plates containing 50 µg/mL of streptomycin. Single colonies were picked and used to inoculate LB supplemented with 50 µg/mL of streptomycin and grown overnight at 37 °C. Target plasmids were isolated using QIAprep Spin Miniprep Kit (Qiagen) and insert sequence was verified by sanger sequencing.

***Expression and purification of ChrA***

The sequence-verified pACYCDuet-His$_6$*chrA* plasmid was used to transform BL21(DE3) electrocompetent cells, plated on Luria Broth (LB) agar containing 33 µg/mL chloramphenicol and incubated at 37 °C overnight. A single colony was used to inoculate 5 mL of LB containing 33 µg/mL chloramphenicol and grown at 37 °C overnight with shaking at 200 rpm. The following day, the overnight starter culture was used to inoculate 4 L of LB containing 33 µg/mL chloramphenicol and grown at 37 °C with shaking at 200 rpm until an O.D$_{600}$ of 0.6 was reached. The culture was cooled on ice for 30 min and ChrA expression was induced by the addition of IPTG to a final concentration of 0.5 mM and grown at 18 °C and 200 rpm for 16 h. The cells were harvested by centrifugation at 6000 *x g* for 10 min at 10 °C. The cell pellet was resuspended in 50 mL of lysis buffer (20 mM $Na_2HPO_4$, 300 mM NaCl, 10 mM imidazole, 6 M guanidine HCl, pH 7.5) per 10 g of cell pellet and stirred for 30 min on ice. The cells were lysed using a high-pressure homogenizer (Avestin, Inc) at an operating pressure between 5000 and 10,000 psi and the lysate was clarified by centrifugation at 18,000 *x g* for 30 min. The supernatant was filtered using a 0.45 µm syringe filter and loaded onto a 10-mL Ni-nitrilotriacetic acid (Ni-NTA) gravity column pre-equilibrated with 2 column volumes (CV) of lysis buffer. The column containing the supernatant and resin was nutated at 4 °C for 30 min before allowing to drain under gravity. The column was then washed with 10 CV of wash buffer 1 (20 mM $Na_2HPO_4$, 300 mM NaCl, 30 mM imidazole, 4 M guanidine HCl, pH 7.5), followed by 10 CV of wash buffer 2 (20 mM $Na_2HPO_4$, 300 mM NaCl, 30 mM imidazole, pH 7.5). ChrA was then eluted using 4 CV of elution buffer (20 mM $Na_2HPO_4$, 300 mM NaCl, 500 mM imidazole, pH 7.5). The eluted peptide was then desalted using Chromabond C8 solid phase extraction cartridges and lyophilized overnight. The lyophilized peptide was redissolved in 5 mL of 5% acetonitrile and 0.1% trifluoroacetic acid (TFA) and purified by reversed-phase HPLC using a Phenomenex C18 column (5 µm, 100 Å, 250 mm ×10 mm) with a 60 min linear gradient from 4% solvent A (0.1% TFA) and 100% solvent B (0.1% TFA in acetonitrile). Peaks containing ChrA were pooled, lyophilized, and stored at -20 $^{o}$C until use.

***Heterologous production of ChrA****

*E. coli* BL21(DE3) electrocompetent cells were transformed with the plasmids pACYCDuet_*His$_6$chrA* and pCDFDuet_*chrH/chrI* and selected on LB agar plates containing 33 µg/mL of chloramphenicol and 50 µg/mL kanamycin and incubated at 37 $^{o}$C overnight. A single colony was used to inoculate 5 mL of LB containing appropriate antibiotics and incubated at 37 $^{o}$C overnight. The culture was used to inoculate 2 L of LB supplemented with 33 µg/mL of chloramphenicol and 50 µg/mL kanamycin and incubated at 37 $^{o}$C with shaking at 200 rpm until an $OD_{600}$ of 0.7 was reached. The culture was cooled on ice for 30 min and IPTG and ferrous ammonium sulfate were added to final concentrations of 0.7 mM and 50 µM respectively. The culture was incubated at 18 $^{o}$C with shaking at 200 rpm for a 18 h. The cells were harvested by centrifugation at 6500 rpm for 10 min at 4 $^{o}$C and the cell pellet was resuspended in 5 x pellet volume of lysis buffer (20 Mm $Na_2HPO_4$, 300 mM NaCl, 6 M guanidine, 10 mM imidazole, pH 7.5) and stirred at room temperature for 30 min. Lysis was completed by passing the lysate through an Avestin high-pressure homogenizer five times with an operating pressure between 5000 and 10,000 psi. The lysate was clarified by centrifugation at 20,000 *x g* for 1 h. The supernatant was loaded onto a 5-mL Ni-NTA gravity column pre-equilibrated with 3 CV of lysis buffer. The column was capped and nutated at 4 $^{o}$C for 30 min before it was mounted and allowed to drain. The column was washed with 10 CV of wash buffer 1 (20 mM $Na_2HPO_4$, 300 mM NaCl, 4 M guanidine, 3 mM imidazole, pH 7.5) followed by 10 CV of wash buffer 2 (20 mM $Na_2HPO_4$, 300 mM NaCl, 30 mM imidazole, pH 7.5). The peptide was then eluted with 3 CV of Elution buffer (20 mM $Na_2HPO_4$, 300 mM NaCl, 500 mM imidazole, pH 7.5). The eluted fractions were desalted with Chromabond C4 Solid Phase Extraction (SPE) cartridges, lyophilized, and stored at -20 $^{o}$C.

***Expression and purification of ChrH***

The plasmid pET28a-His*6chrH* was used to transform Rosetta(DE3)pLysS electrocompetent cells, plated on Luria Broth (LB) agar containing 50 µg/mL of kanamycin and 33 µg/mL chloramphenicol and incubated at 37 $^{o}$C overnight. A single colony was used to inoculate 5 mL of LB containing 50 µg/mL of kanamycin and 33 µg/mL chloramphenicol and grown at 37 $^{o}$C overnight with shaking at 200 rpm. The following day, the overnight starter culture was used to inoculate 4 L of LB containing 50 µg/mL of kanamycin and 33 µg/mL chloramphenicol and grown at 37 $^{o}$C with shaking at 200 rpm until an $O.D_{600}$ of 0.6. The culture was cooled on ice for 30 min followed by the addition of ferrous citrate and IPTG to final concentrations of 250 µM and 0.5 mM respectively and grown for an additional 16 h at 18 $^{o}$C and 200 rpm. The cells were pelleted by centrifugation at 6000 *x g* for 10 min at 10 $^{o}$C. The cell pellet was resuspended in 50 mL of lysis buffer (20 MOPS, 300 mM NaCl, 30 mM imidazole, 10% glycerol, pH 7.5) per 10 g of cell mass. To this suspension were added 50 mg of lysozyme, 50 mg of CHAPS, 10 µL/mL of Protease Inhibitor Cocktail (Thermo Scientific), and 50 µL of DNAse I (1000 U/mL) per liter of cell culture and stirred for 30 min on ice. The lysate was clarified by centrifugation at 18,000 *x g* for 30 min. The supernatant was filtered using a 0.45 µm syringe filter and loaded onto a 10-mL Ni-NTA gravity column pre-equilibrated with 2 CV of lysis buffer. The resin-lysate mixture was rocked for 10 min and allowed to drain under gravity. The resin was washed with 10 CV of Lysis buffer and the bound protein was eluted using elution buffer (20 MOPS, 300 mM NaCl, 300 mM imidazole, 10% glycerol, pH 8.0). Eluted fractions were visualized on 4-20% SDS-PAGE (Bio-Rad) and ChrH-containing fractions were pooled and desalted using a PD-10 column (Cytiva) according to the manufacturer's protocol into storage buffer (20 MOPS, 300 mM NaCl, 10% glycerol, pH

7.5). The flow-through was concentrated using 30 kDa cutoff Amicon Ultra-15 centrifugal filter unit (Millipore). The concentration of ChrH was estimated using nanodrop.

***Analysis of metal concentration***

Iron quantification in ChrH was carried out according to a previously published protocol.[1,2] Briefly, ChrH was purified by size-exclusion chromatography (SEC) using a Superdex 200 column. The protein was eluted with Tris buffer (20 mM Tris, 300 mM NaCl, 10% glycerol, pH 7.6) at 1 mL/min flow rate. ChrH was then reconstituted with 10 mM $FeSO_4$, 20 mM ascorbic acid in 20 mM Tris, pH 7.6. Excess iron was removed from ChrH using a PD10 column prior to iron analysis. Iron quantification of ChrH was determined using Ferene S as a spectrophotometric dye according to the protocol reported by Hennessy and co-workers.[3] A standard curve was generated using an iron standard in 2% $HNO_3$ solution (Claritas PPT).

***Expression and purification of ChrI***

*E. coli* RosettapLysS electrocompetent cells were transformed with the plasmids pET28a_His$_6$*ChrI* and selected on agar plates supplemented with 33 µg/mL of chloramphenicol and 50 µg/mL of kanamycin. A single colony was used to inoculate 5 mL of LB supplemented with appropriate antibiotics and incubated at 37 $^{o}$C overnight with shaking. The overnight preculture was used to inoculate 2 L of TB containing 33 µg/mL of chloramphenicol and 100 µg/mL of kanamycin and incubated at 37 $^{o}$C with shaking at 220 rpm until an $OD_{600}$ of 1.2 was reached. The culture was cooled on ice for 30 min and expression was induced by addition of IPTG to a final concentration of 0.7 mM. The culture was incubated at 18 $^{o}$C and 220 rpm for an additional 18 h before the cells were harvested by centrifugation at 6500 *x g* for 10 min at 4 $^{o}$C. A total of 30 g of cell pellet was obtained which was resuspended in 150 mL of lysis buffer (20 mM MOPS, 300 mM NaCl, 30 mM imidazole, 10% glycerol, pH 7.5). To the suspension were added 50 mg of lysozyme, 50 mg of CHAPS, 10 µL/mL of Protease Inhibitor Cocktail, and 50 µL of DNAse I (1000 U/mL) per liter of cell culture and stirred for 30 min on ice. Lysis was completed by passing the resulting mixture through an Avestin Inc. high-pressure homogenizer 5 times with an operating pressure between 5000 and 10000 psi. To the lysate was added Triton X-114 (Sigma) to a final concentration of 1% and stirred at 4 $^{o}$C for 16 h. The lysate was clarified by centrifugation at 20,000 *x g* for 1 h at 4 $^{o}$C. The supernatant was loaded onto a 5-mL Ni-NTA gravity column pre-equilibrated with lysis buffer. The column was capped and nutated at 4 $^{o}$C for 30 min after which the column was drained under gravity. The resin was washed with 10 CV of lysis buffer containing 0.01% of n-dodecyl-β-maltoside (DDM). Bound proteins were eluted with elution buffer (20 mM MOPS, 300 mM NaCl, 30 mM imidazole, 10% glycerol, pH 7.5, 0.01% DDM). Eluted fractions containing ChrI were pooled, and buffer exchanged into storage buffer. The purity of the protein was assessed by 4-20 % SDS PAGE, flash-frozen in liquid nitrogen and stored at -80 $^{o}$C until use.

***In vitro reconstitution of ChrHI activity***

In vitro assays were carried out in 1.5 mL Eppendorf tubes on a 100 µL scale and contained 20 mM MOPS, 300 mM NaCl, pH 7.5 as the reaction buffer. Consecutively, 200 µM ChrA, 2 mM dithiothreitol (DTT), 100 µM ChrI, 100 µM ChrH and 2 mM SAM were added and incubated at room temperature overnight. Control reactions were set up in a similar manner except buffer was substituted for reagents, protein, or substrate. The reactions were quenched by adding 10 µL of 10 x quenching buffer (100 mM sodium citrate, 0.5 mM EDTA, pH 3.0) and centrifuged at 6000 *x g* for 5 min to remove precipitates. Detergents used in ChrI purification were subsequently removed using ThermoFisher detergent removal spin columns according to the manufacturer's protocol. The resulting flow-through fractions were ziptipped using Agilent C18 ziptips and analyzed by MALDI-TOF MS for modifications. ChrA and modified ChrA were digested with endoproteinase Lys at 37 $^{o}$C overnight and purified on a reversed-phase Phenomenex C8 column using an Agilent HPLC system with 0.1% TFA and acetonitrile as the mobile phase. Peaks corresponding to linear and modified peptides were collected, lyophilized, and analyzed by MALDI-TOF MS and HRMS.

##### *Labeling of ChrA* with of $^{13}CD_3$-Methionine isotope and Selenomethionine.*

Incorporation of $^{13}CD_3$-methionine and selenomethionine into ChrA* was carried out according to a previously published protocol.[4] Briefly, the plasmids pACYCDuet_His$_6$*chrA* and pCDFDuet_*chrH/chrI* were used to co-transform *E. coli* BL21 (DE3) and transformants were selected on LB agar plates supplemented with 33 µg/mL chloramphenicol and 50 µg/mL streptomycin. A single colony was used to inoculate 5 mL of LB containing appropriate antibiotics and grown at 37 $^{o}$C overnight. The overnight culture was centrifuged for 10 min at 4500 *x g* and the pellet was used to inoculate 250 mL of M9 minimal medium containing 30 mM $KH_2PO_4$, 23 mM $K_2HPO_4$, 16 mM $Na_2HPO_4$, 17 mM NaCl, 37 mM $NH_4Cl$, pH 7.4 supplemented with trace metal stock solution (Teknova), 2 mM $MgSO_4$, 0.4% w/v glucose, 50 mg/L thiamine and 33 µg/mL chloramphenicol and 50 µg/mL streptomycin. The defined medium preculture was incubated at 37 $^{o}$C overnight with shaking at 200 rpm. The following day, the cell density reached an $OD_{600}$ of approximately 1.5. The cells were harvested by centrifugation 5000 *x g* for 10 min and pellet was resuspended in 250 mL of fresh M9 medium of which 20 mL was used to inoculate 1 L of the same M9 medium. The fresh culture was incubated at 37 $^{o}$C with shaking at 200 rpm until an $OD_{600}$ of approximately 0.6 was reached. The culture was cooled on ice for 30 min at which point 100 mg L-$^{13}CD_3$-methionine (Cambridge Stable Isotopes) or L-selenomethionine (TCI chemicals) was added along with 100 mg each of lysine, threonine, phenylalanine, and 50 mg each of leucine, isoleucine, and valine. The expression culture was incubated at 18 $^{o}$C for 30 min before IPTG and iron(II)citrate were added to final concentrations of 0.7 mM and 25 µM, respectively. Expression was continued for an additional 18 h at 18 $^{o}$C and 220 rpm. Cells were harvested by centrifugation at 6500 *x g* for 10 min at 4 $^{o}$C and purified as described above.

##### *Labeling of ChrA* with 1-$^{13}C$-Cysteine*

The *E. coli* strain JW3582-2, a cysteine auxotroph, was lysogenized using λDE3 Lysogenization Kit (Novagen) to allow for expression of target genes in pET vectors. The plasmids pACYCDuet-*chrA* and pCDFDuet-*chrHI* were used to transform the electrocompetent cells and transformants were selected on LB agar plates supplemented with 50 µg/mL of kanamycin, 33 µg/mL of chloramphenicol and 50 µg/mL of streptomycin. A single colony was used to inoculate 5 mL of LB supplemented with appropriate antibiotics and grown at 37 $^{o}$C overnight. The preculture was harvested by centrifugation at 5000 *x g* for 10 min and the pellet was used to inoculate 1 L of defined medium (22 mM $KH_2PO_4$, 33.7 mM $Na_2HPO_4$, 8.6 mM NaCl, pH 7.4) supplemented with 3 mL of 40% $(NH_4)_2SO_4$, 100 µL of $CaCl_2$, 2 mL of $MgSO_4$, 1 mL of 1000x biotin (1 mg/mL), 1 mL of 1000x Trace metals (1.25 mM $ZnCl_2$, 260 µM $CuCl_2.H_2O$, 252 µM $CoCl_2.6H_2O$, 250 µM $Na_2MoO_4.2H_2O$, pH 7.4), 20 mL of 50x MEM amino acid solution (Thermofisher), 50 µg/mL of kanamycin, 33 µg/mL of chloramphenicol and 50 µg/mL of streptomycin and incubated at 37 $^{o}$C with shaking at 200 rpm for additional 16 h. The culture was centrifuged at 5000 *x g* and the pellet was washed with 500 mL of phosphate buffer (22 mM $KH_2PO_4$, 33.7 mM $Na_2HPO_4$, 8.6 mM NaCl, pH 7.4), resuspended in 200 mL of fresh phosphate buffer and 50 mL was used to inoculate 1 L of fresh defined medium supplemented with 3 mL of 40% $(NH_4)_2SO_4$, 100 µL of $CaCl_2$, 2 mL of $MgSO_4$, 1 mL of 1000x biotin (1 mg/mL), 1 mL of 1000x trace metal solution (1.25 mM $ZnCl_2$, 260 µM $CuCl_2.H_2O$, 252 µM

$CoCl_2.6H_2O$, 250 µM $Na_2MoO_4.2H_2O$, pH 7.4), 20 mL of 50x Δ L-cysteine MEM amino acid solution (30 mM L-arginine hydrochloride, 10 mM L-histidine hydrochloride, 20 mM L-isoleucine, 20 mM L-leucine, 20 mM L-lysine hydrochloride, 5 mM L-methionine, 10 mM L-phenylalanine, 20 mM L-threonine, 2.5 mM L-tryptophan, 10 mM L-tyrosine and 20 mM L-valine), 50 µg/mL of kanamycin, 33 µg/mL of chloramphenicol, 50 µg/mL of streptomycin, and 100 mg of 1-$^{13}$C-L-cysteine and incubated at 37 $^{o}$C with shaking at 220 rpm until an $OD_{600}$ of 0.7 was reached. The culture was cooled on ice for 30 min and incubated at 18 $^{o}$C for 30 min before IPTG and iron(II) citrate were added to final concentrations of 0.7 mM and 25 µM, respectively. The culture was incubated for an additional 18 h at 18 $^{o}$C with shaking at 220 rpm. The cells were harvested by centrifugation at 6500 *x g* at 4 $^{o}$C for 10 min and modified ChrA was purified as described above.

##### *Preparation of S-adenosyl-L-methionine synthetase (MetK)*

Purification of SAM synthetase (MetK) from *Bacillus subtilis* was carried out as previously described.[5-7] Briefly, the codon-optimized synthetic gene encoding *Bacillus subtilis* SAM synthetase ( MetK, Uniprot ID P54419) I317V variant inserted between the NdeI and XhoI restriction sites of the pET28a was purchased from Twist Biosciences (South Francisco, CA). The MetK[I317V]- pET28a vector was used to transform *E. coli* BL21(DE3) electrocompetent cells. A single colony was used to inoculate 5 mL of LB supplemented with 50 µg/mL of kanamycin and grown at 37 $^{o}$C overnight. The overnight culture was used to inoculate 2 x 1 L of LB containing 50 µg/mL of kanamycin and grown at 37 $^{o}$C with shaking until an 0D600 of 0.6 was reached. The culture was cooled on ice for 30 min and protein expression was induced with 0.5 mM IPTG after which the cell culture was incubated at 18 $^{o}$C for an additional 18 h with shaking at 200 rpm. The cells were harvested by centrifugation at 6500 *x g* for 10 min and the cell pellet (15 g) was resuspended in 100 mL of lysis buffer (20 mM Tris HCl, 300 mM NaCl, 40 mM imidazole, 10% glycerol, pH 8.0). To the suspension was added 50 mg of Lysozyme, 50 mg of CHAPS, 10 µL/mL of Protease Inhibitor Cocktail, and 50 µL of DNAse I (1000 U/mL) per liter of cell culture and stirred for 30 min on ice. The lysate was passed through a high-pressure homogenizer (Avestin Inc) four times with an operating pressure between 5,000 and 10,000 psi followed by centrifugation at 15,000 *x g* for 30 min. The supernatant was loaded onto 5 mL of Ni-NTA gravity column pre-equilibrated with lysis buffer. The column was capped and nutated at 4 $^{o}$C for 30 min and allowed to drain under gravity. The column was washed with 5 CV of lysis buffer and bound proteins were eluted with 2 CV of elution buffer (20 mM Tris HCl, 300 mM NaCl, 300 mM imidazole, 10% glycerol, pH 8.0). Elution fractions were visualized on SDS-PAGE and fractions containing MetK SAM synthetase were pooled and concentrated to 2.5 mL using 10 kDa Amicon spin concentrators (Sigma). Proteins were exchanged into storage buffer (20 mM Tris HCl, 300 mM NaCl, 10% glycerol, pH 8.0) using PD-10 columns, concentrations were estimated using a Nanodrop, aliquoted into 100 µL volumes, flash-frozen in liquid nitrogen and store at -80 $^{o}$C until use.

##### *Enzymatic synthesis of S-adenosyl-L-methionine (SAM) and its isotopologs*

To a gently stirred solution of 10 mM methionine, 100 mM Tris HCl and 50 mM KCl in 16.2 mL of $ddH_2O$ were added ATP and $MgCl_2$ to final concentrations of 13 mM and 26 mM, respectively. After the solution became homogenous, 4.8 mL of acetonitrile was added and stirring was continued for an additional 30 min. The reaction was initiated by the addition of 100 µM MetK[I317V] and gently stirred at room temperature for 5 h. The resulting mixture was diluted with 80 mL of cold $ddH_2O$ and loaded onto a column packed with 10 mL of Dowex®50WX8 (hydrogen form) that had been pre-equilibrated with $ddH_2O$. The column was washed with 100 mL of 0.5 M cold HCl and eluted with 6 M HCl. The eluted fractions were checked on a Nanodrop and fractions with absorbance above 0.15 at 260 nm were pooled and concentrated with a rotary evaporator. The resulting residue was redissolved in 1 mL of ddH2O and precipitated with 30 mL of acetone. The supernatant was decanted, and the pellet was lyophilized overnight. The solid was redissolved in dd$H_2O$ and the purity was accessed by HR-MS. Likewise, $^{13}CD_3$-SAM was synthesized with $^{13}CD_3$-methionine as starting material.

***Generation of a sequence similarity network of DUF692***

The EFI-EST online server was used to generate a sequence similarity network (SSN)[8] using the families input option. The input family used was PF05114 (DUF692) with a total of 13,035 input sequences (as of November 2, 2022). Default settings were used for all the options for the first step of SSN generation except the SSN edge calculation option for which an E-value of 50 was used. For the initial calculation, an alignment score of 80% was used and the data generated was submitted for analysis using the EFI-Genome Neighborhood Tool. A neighborhood size of 10 and a minimal co-occurrence percentage limit of 20 was used. The final SSN was visualized in cytoscape using the yFiles Organic Layout.

***NMR acquisition***

Samples were dissolved in 280 μL of DMSO-$d_6$ (99.9 atom %, Cambridge Isotope Laboratories) and transferred into a DMSO-matched Shigemi NMR tube, or in 90% $H_2O$ and 10% $D_2O$ in a $D_2O$-matched Shigemi NMR tube. All one and two-dimensional NMR data including $^1H$, $^{13}C$, $^{15}N$, $^1H$-$^1H$ dqCOSY, $^1H$-$^1H$ TOCSY, $^1H$-$^1H$ NOESY, $^1H$-$^{13}C$ HSQC and HMBC, and $^1H$-$^{15}N$ HSQC and HMBC were collected at 23 $^{o}C$ on a Bruker Avance NEO 600 MHz spectrometer with a 5-mm prodigy BBO probe using Topspin 4.1.4 software, or on an Agilent VNMRS 750 MHz spectrometer with a 5-mm HCN probe with VNMRJ 4.2.A software. Typical acquisition parameters included: 1.2 s relaxation delay, 32–64 scans, 16 dummy scans, 2048×256–400 data points, 10 ppm $^1H$ spectral width (in F2 and F1 for homonuclear experiments), or 10 ppm and up to 220 ppm in $^{13}C$ (in F2 and F1), or up to 400 ppm in $^{15}N$ (F1), and 80 ms and 400 ms, or 500 ms mixing time for TOCSY and NOESY, respectively. The dpfgse (double pulsed field gradient spin echo) pulse sequence was used for water suppression (if required). All spectra were processed in MestReNova 14.3.0 and chemical shift assignments based on all 1D and 2D data are shown in Supplementary Tables 1 and 2, respectively.

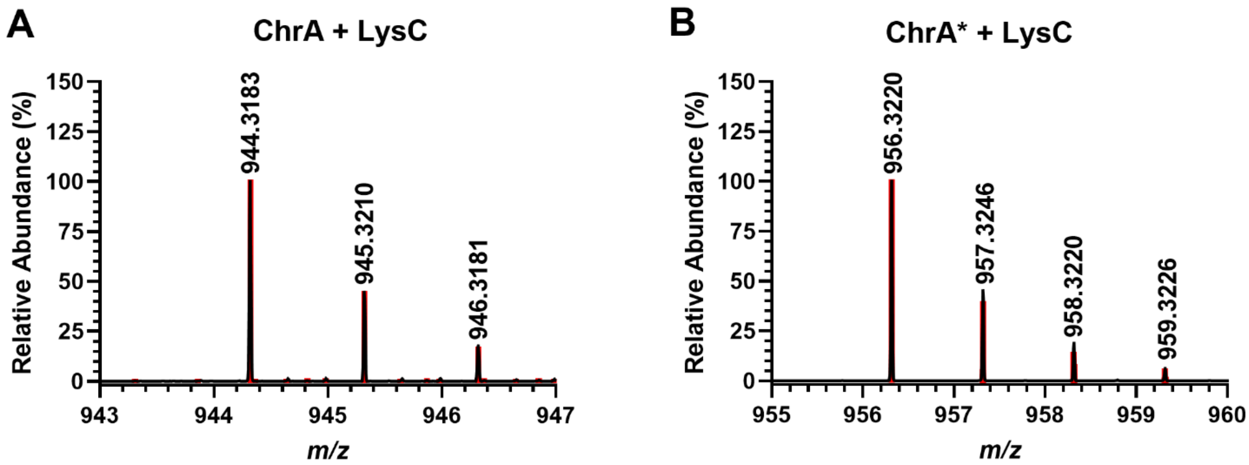

**Figure S1.** High-resolution electrospray ionization mass spectrometry (HRMS) analysis of LysC-digested ChrA and ChrA*. Shown are spectra for **(A)** C-terminal 11mer of ChrA, $[M+H]^+$ calculated $m/z$ = 944.3172, observed $m/z$ = 944.3183 (1.16 ppm error) and **(B)** C-terminal 11mer of ChrA*, $[M+H]^+$ calculated $m/z$ = 956.3172, observed $m/z$ = 956.3220 (5.01 ppm error). Predicted isotope distribution is indicated in red.

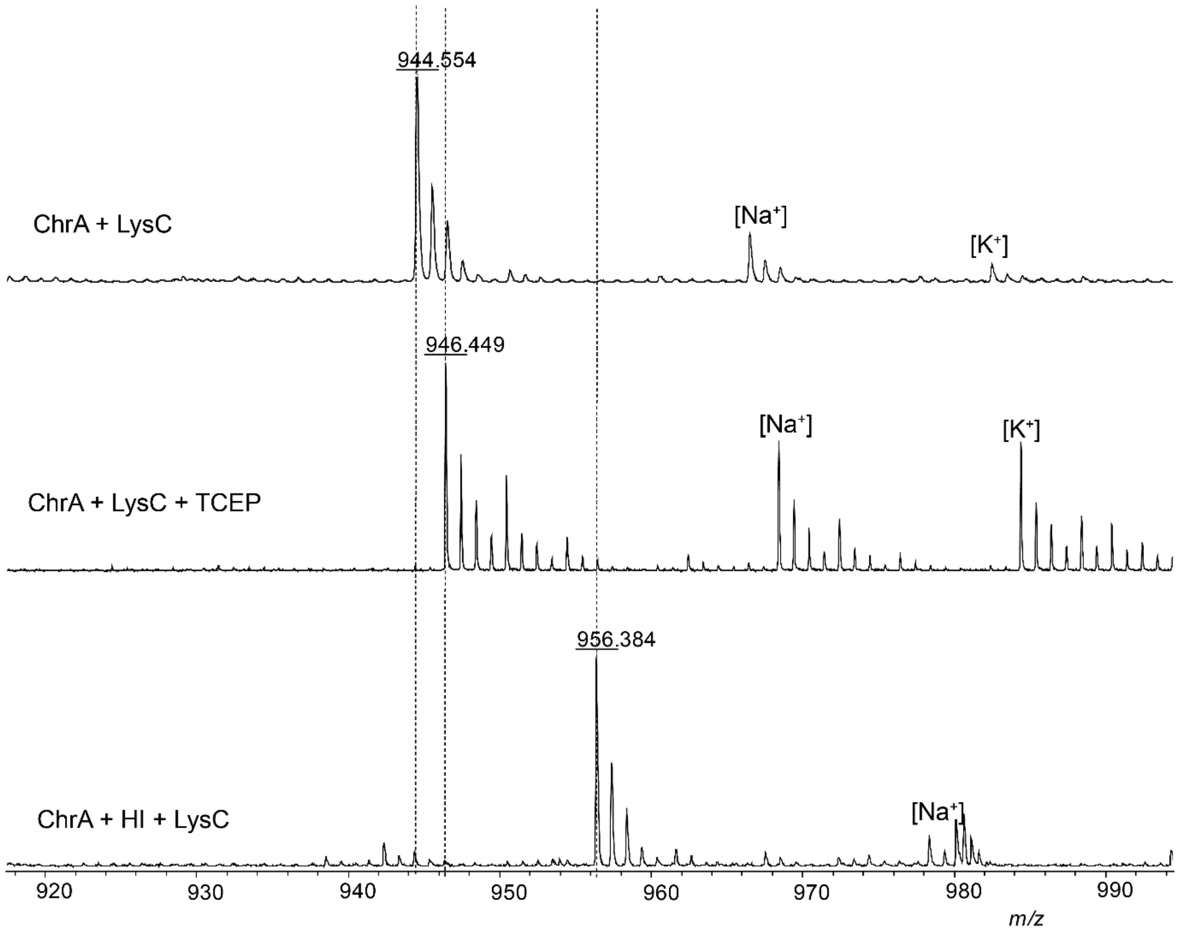

**Figure S2.** Characterization of LysC-digested ChrA and ChrA*. Shown are MALDI-TOF mass spectra of ChrA containing a disulfide bond (top; calc'd *m/z* 944.3172), ChrA treated with TCEP (middle), and ChrA* (bottom). The dashed lines indicate the expected *m/z* of modified and unmodified peptides.

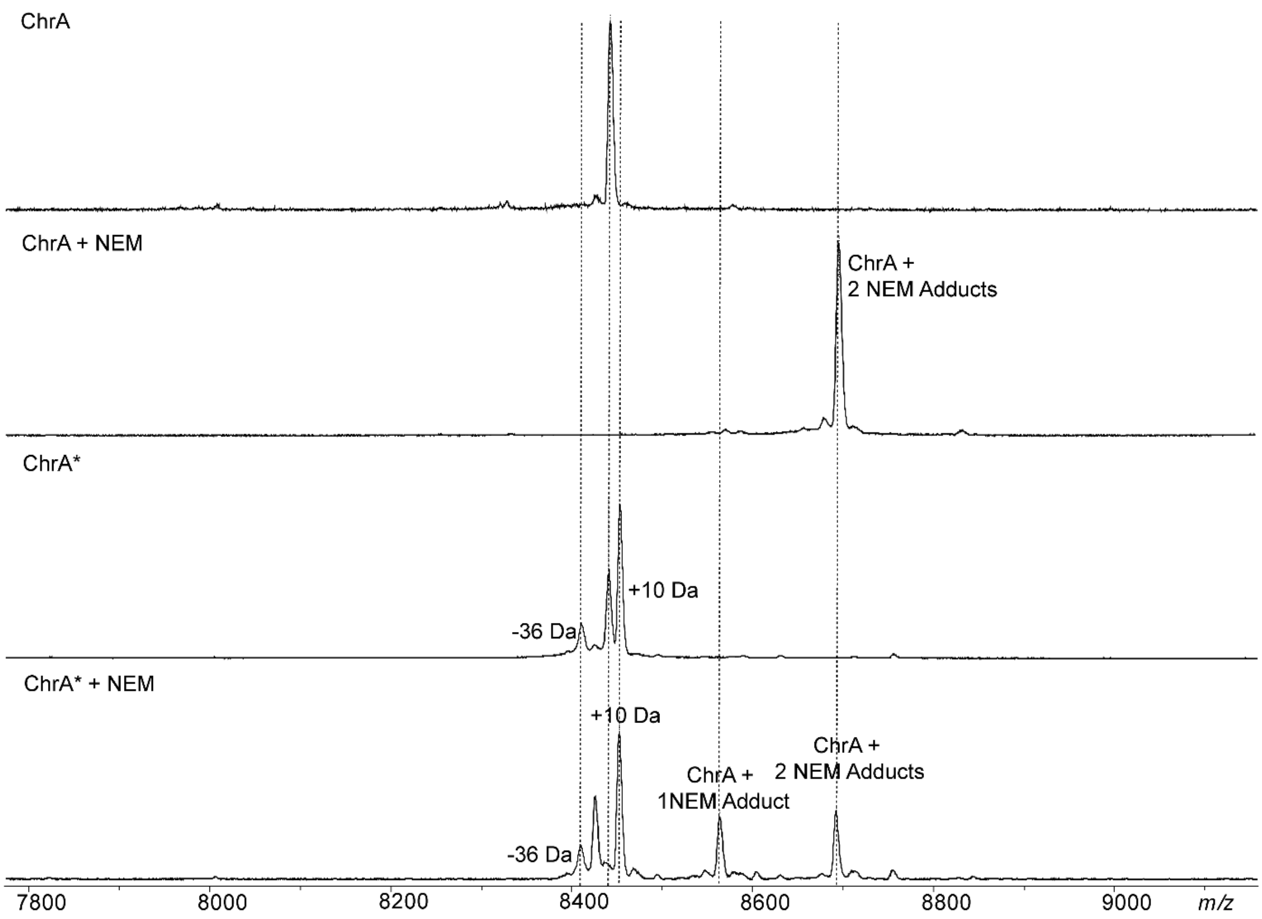

**Figure S3.** Characterization of modified ChrA. Shown are the MALDI-TOF mass spectra of ChrA, ChrA derivatized with NEM, ChrA modified by ChrHI, and modified ChrA derivatized with NEM. The dashed lines indicate the expected *m/z* of modified and unmodified peptides and NEM adducts.

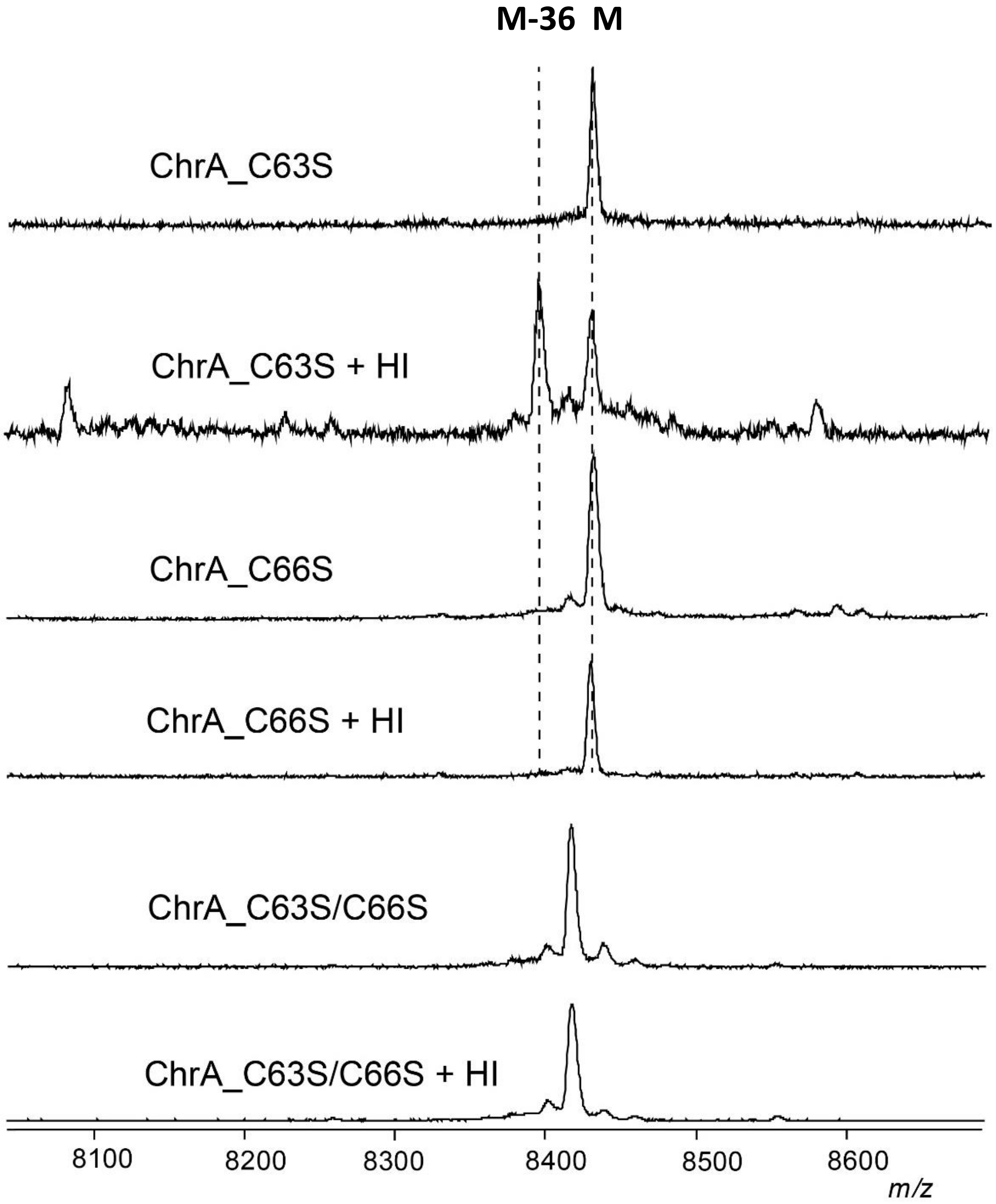

**Figure S4.** Mutational analysis of ChrA. Shown are the MALDI-TOF mass spectra of C66S and C63S mutants of ChrA co-expressed with ChrHI. The dashed lines indicate the expected *m/z* of unmodified peptides and the -36 Da species.

**A**

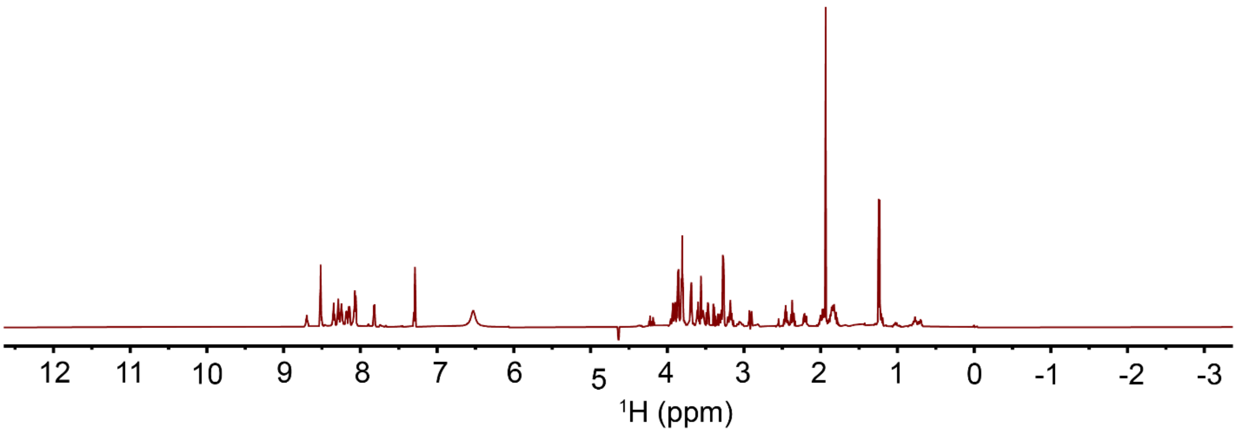
12
11
10
9
8
7
6
5
4
3
2
1
0
-1
-2
-3
$^1$H (ppm)

**B**

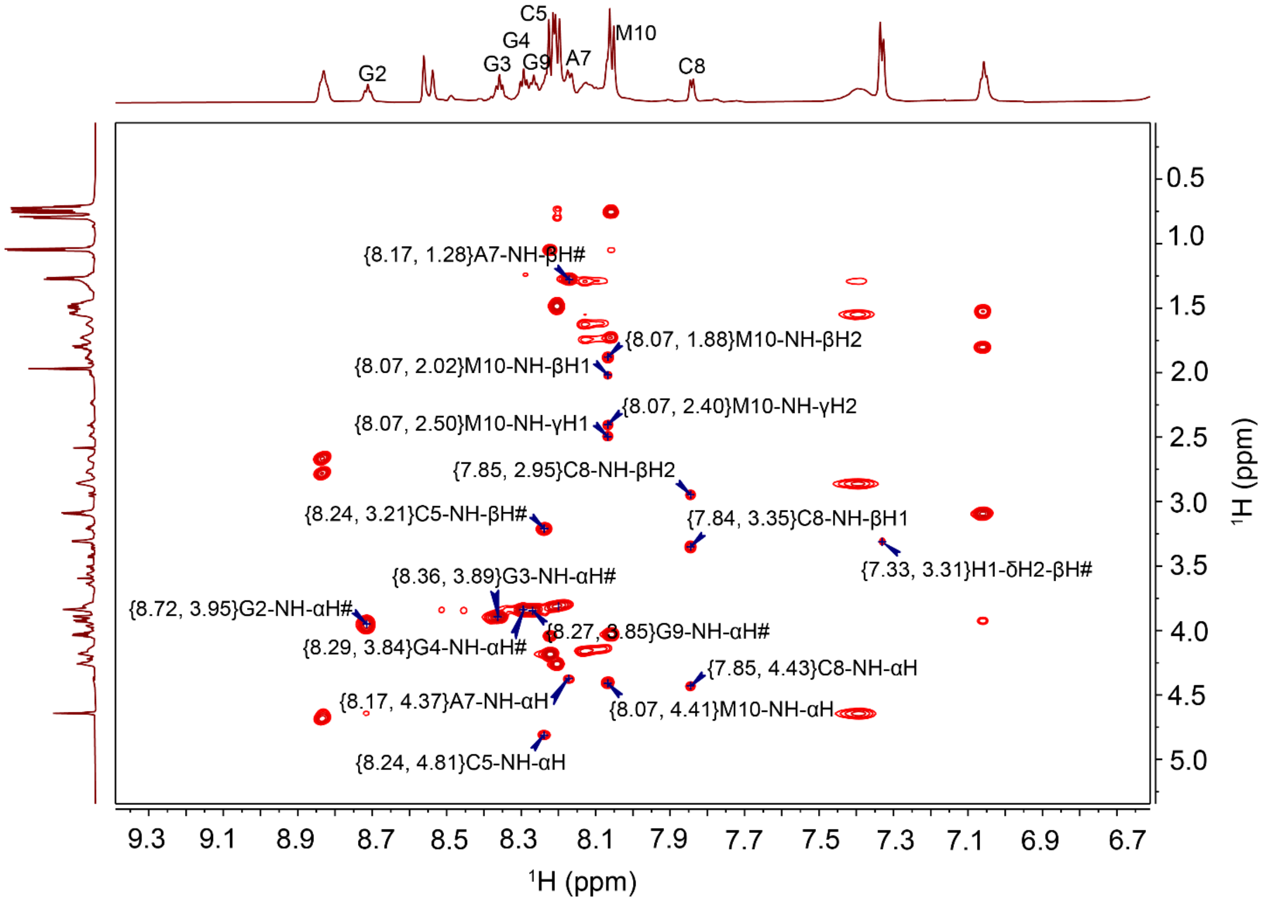
G2
G3
G4
C5
G9
A7
M10
C8
{8.17, 1.28}A7-NH-βH#
{8.07, 1.88}M10-NH-βH2
{8.07, 2.02}M10-NH-βH1
{8.07, 2.40}M10-NH-γH2
{8.07, 2.50}M10-NH-γH1
{7.85, 2.95}C8-NH-βH2
{8.24, 3.21}C5-NH-βH#
{7.84, 3.35}C8-NH-βH1
{7.33, 3.31}H1-δH2-βH#
{8.36, 3.89}G3-NH-αH#
{8.72, 3.95}G2-NH-αH#
{8.27, 3.85}G9-NH-αH#
{8.29, 3.84}G4-NH-αH#
{7.85, 4.43}C8-NH-αH
{8.17, 4.37}A7-NH-αH
{8.07, 4.41}M10-NH-αH
{8.24, 4.81}C5-NH-αH
9.3
9.1
8.9
8.7
8.5
8.3
8.1
7.9
7.7
7.5
7.3
7.1
6.9
6.7
$^1$H (ppm)
0.5
1.0
1.5
2.0
2.5
3.0
3.5
4.0
4.5
5.0
$^1$H (ppm)

C

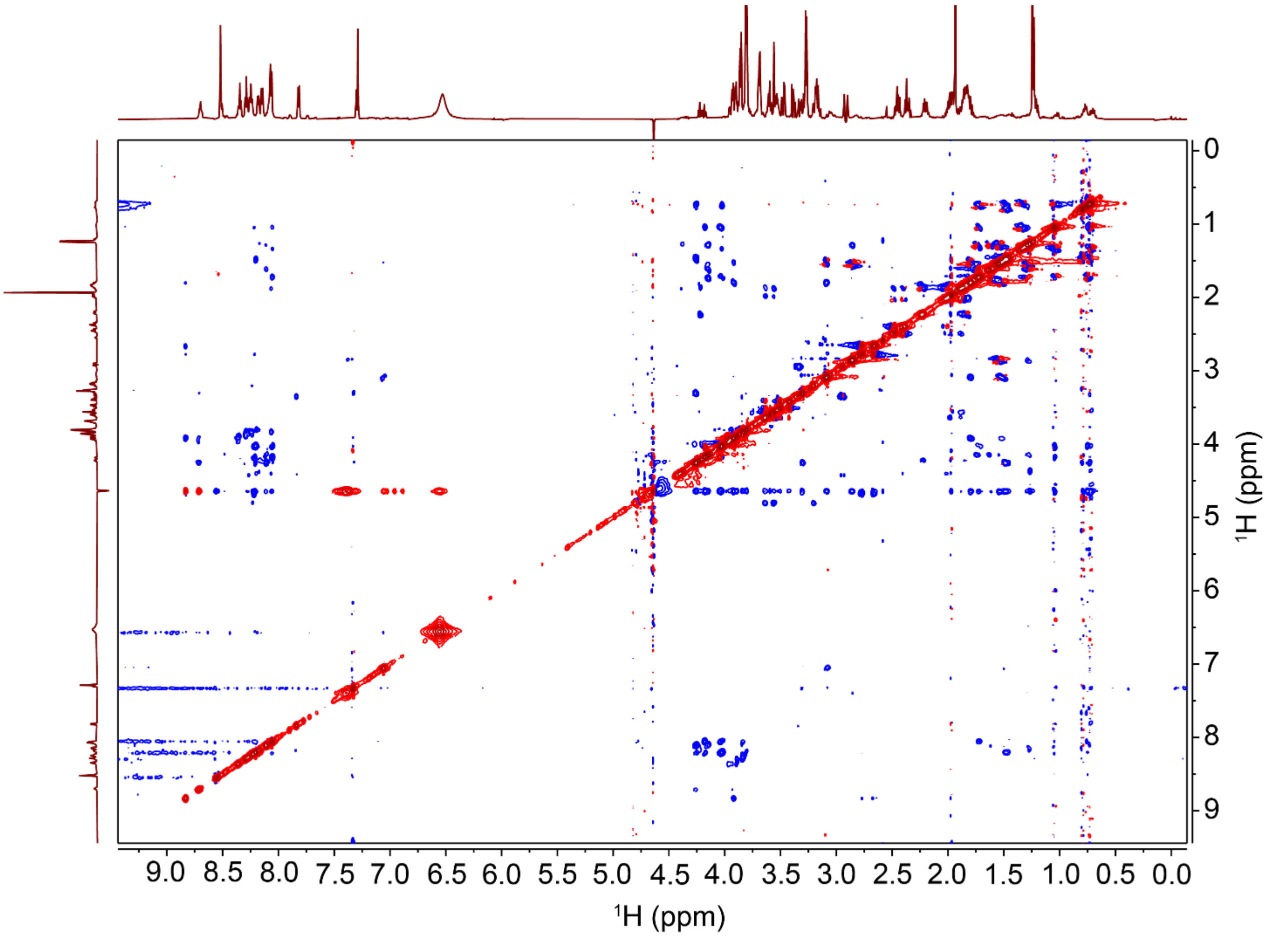

9.0 8.5 8.0 7.5 7.0 6.5 6.0 5.5 5.0 4.5 4.0 3.5 3.0 2.5 2.0 1.5 1.0 0.5 0.0
$^1$H (ppm)
0 1 2 3 4 5 6 7 8 9
$^1$H (ppm)

**D**

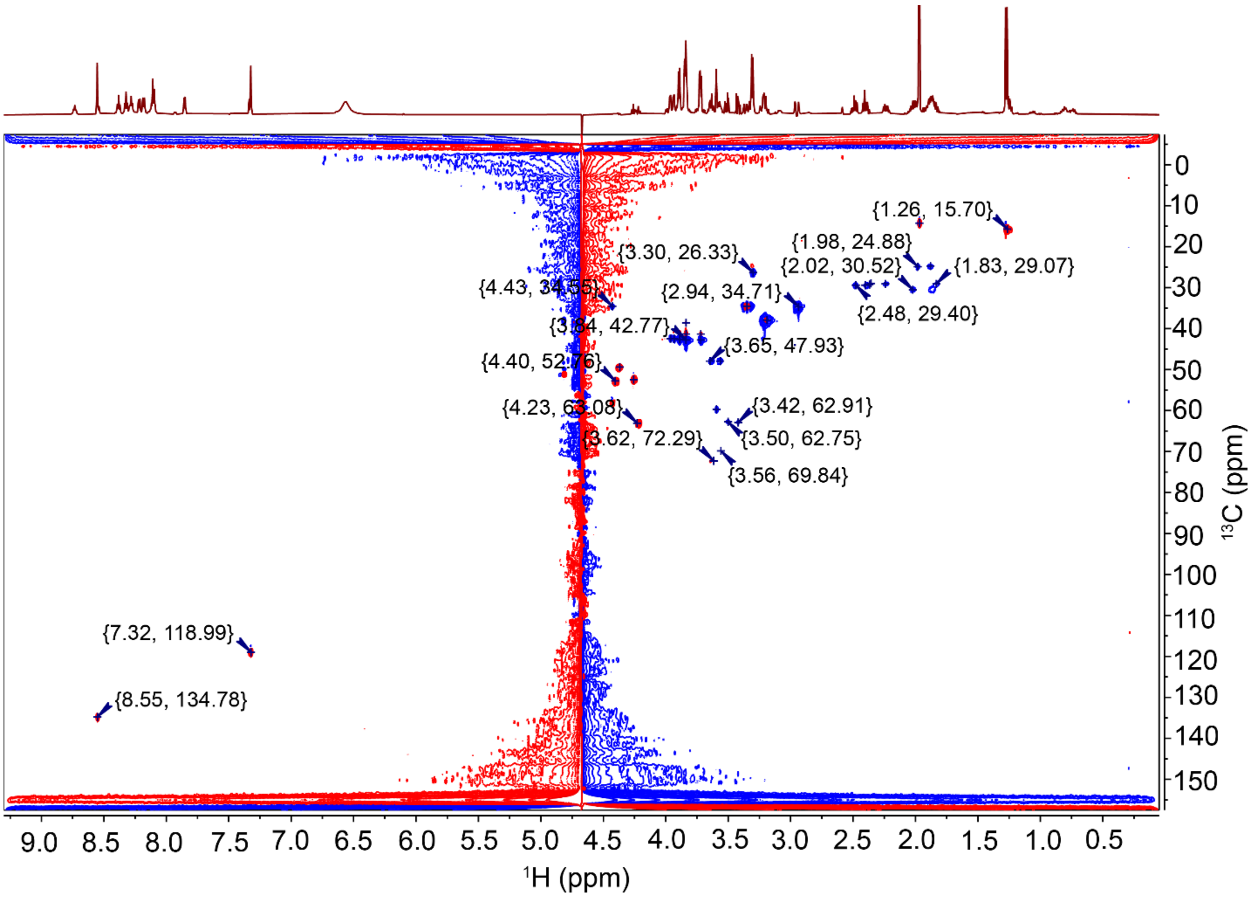

**Figure S5.** NMR spectra of LysC-cleaved ChrA peptide in 90% $H_2O$ and 10% $D_2O$ at 25 $^{o}C$. Shown are (A) $^{1}H$, (B) TOCSY, (C) ROESY and (D) $^{1}H$-$^{13}C$ HSQC spectra.

**A**

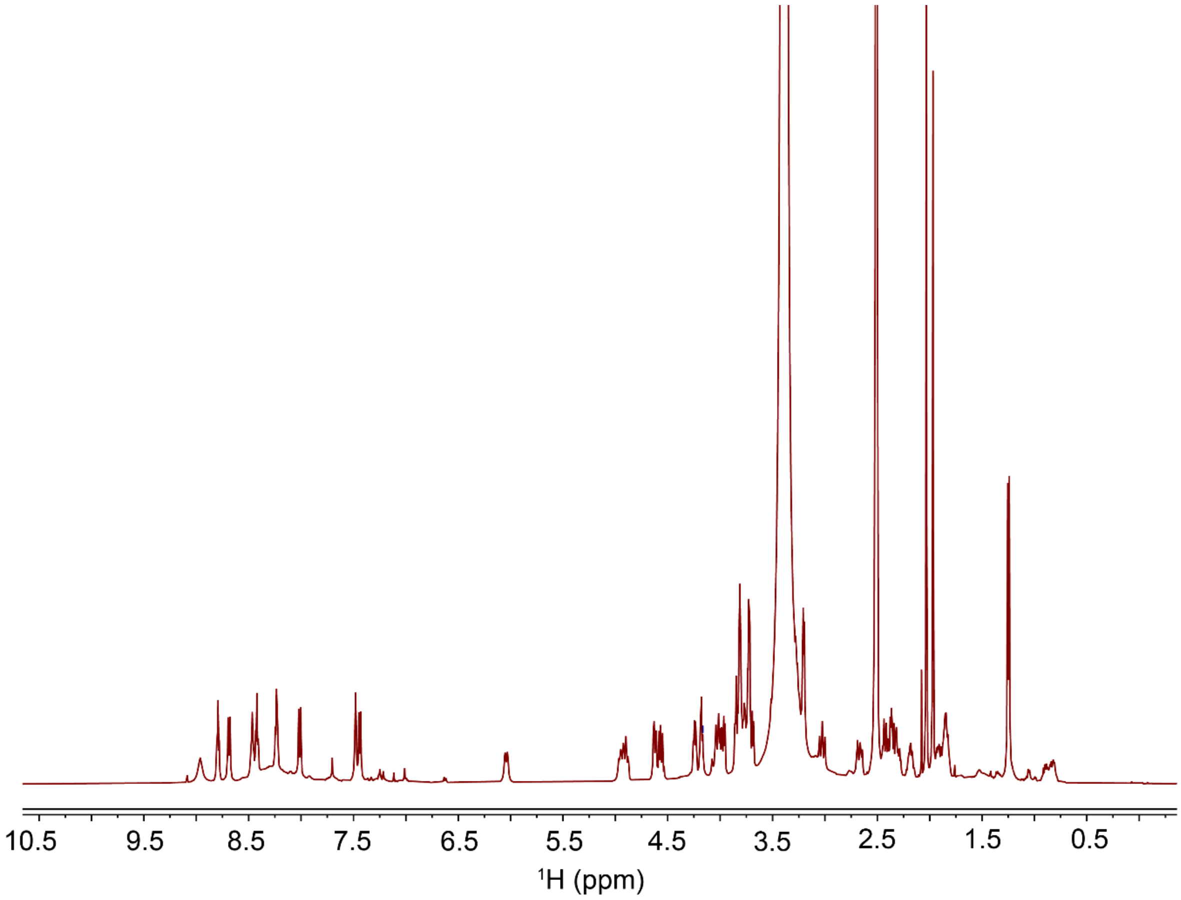

**B**

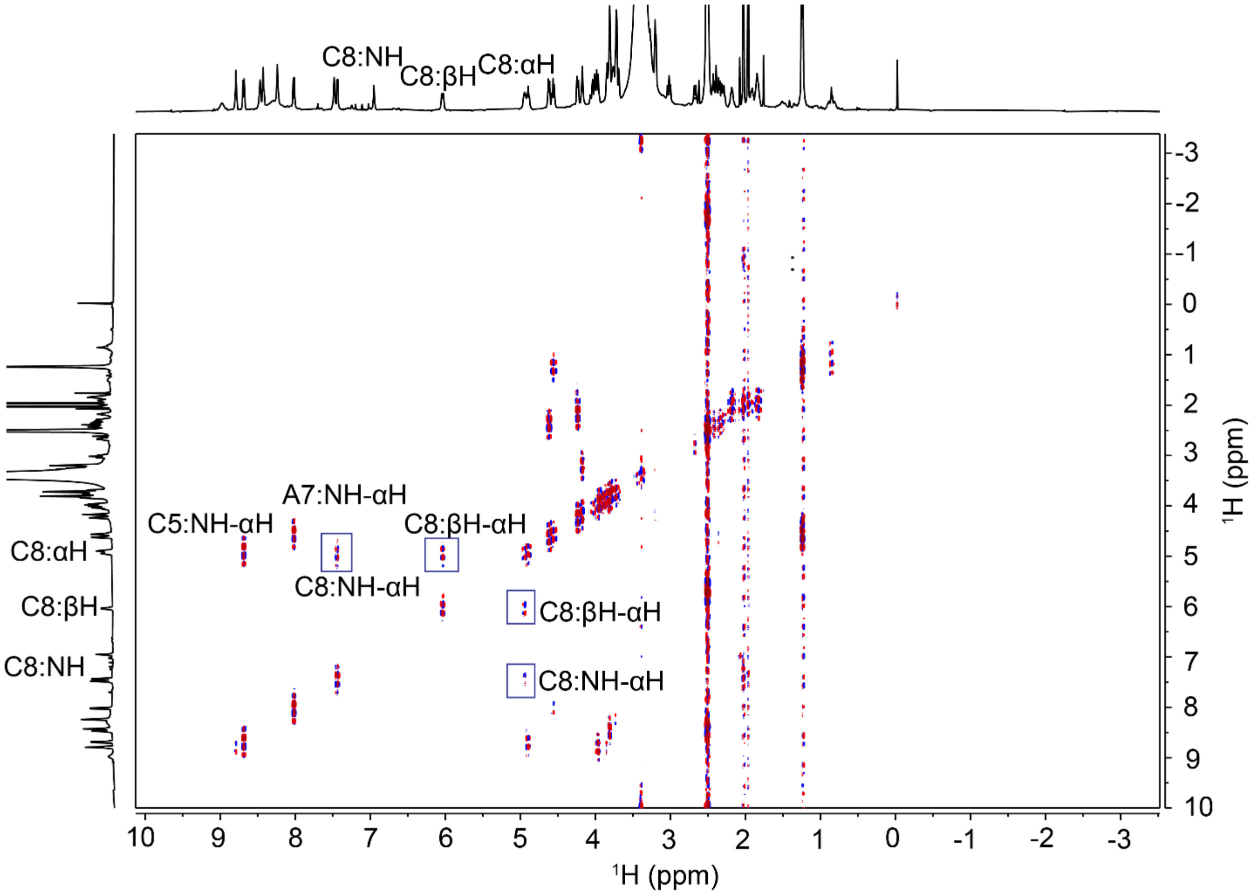
C8:NH
C8:βH
C8:αH
A7:NH-αH
C5:NH-αH
C8:βH-αH
C8:NH-αH
C8:βH-αH
C8:NH-αH
C8:αH
C8:βH
C8:NH
$^1$H (ppm)
$^1$H (ppm)

**C**

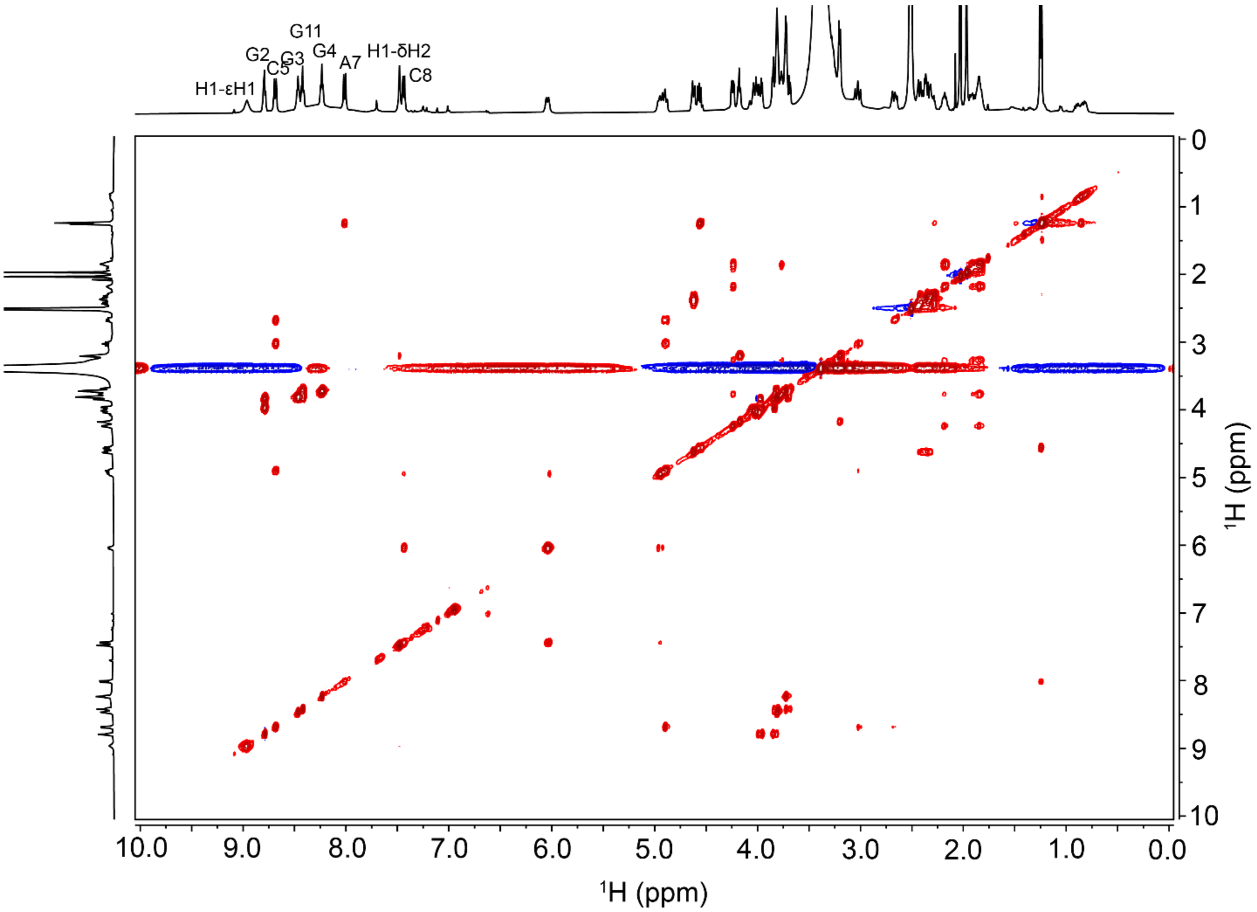

**D**

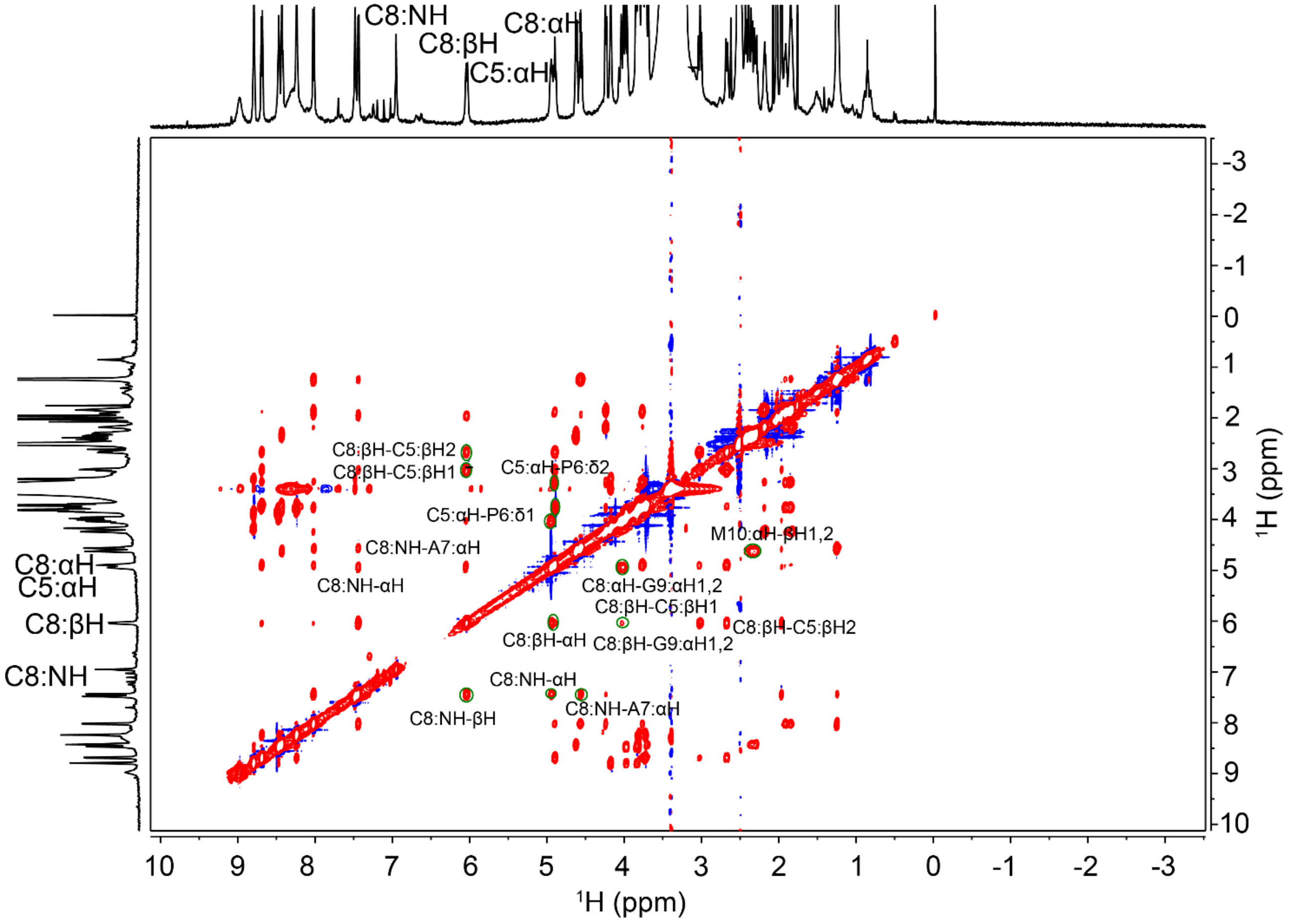
C8:NH
C8:βH
C8:αH
C5:αH
C8:βH-C5:βH2
C8:βH-C5:βH1
C5:αH-P6:δ2
C5:αH-P6:δ1
C8:NH-A7:αH
M10:αH-βH1,2
C8:NH-αH
C8:αH-G9:αH1,2
C8:βH-C5:βH1
C8:βH-C5:βH2
C8:βH-αH
C8:βH-G9:αH1,2
C8:NH-αH
C8:NH-A7:αH
C8:NH-βH
$^1$H (ppm)
$^1$H (ppm)

**E**

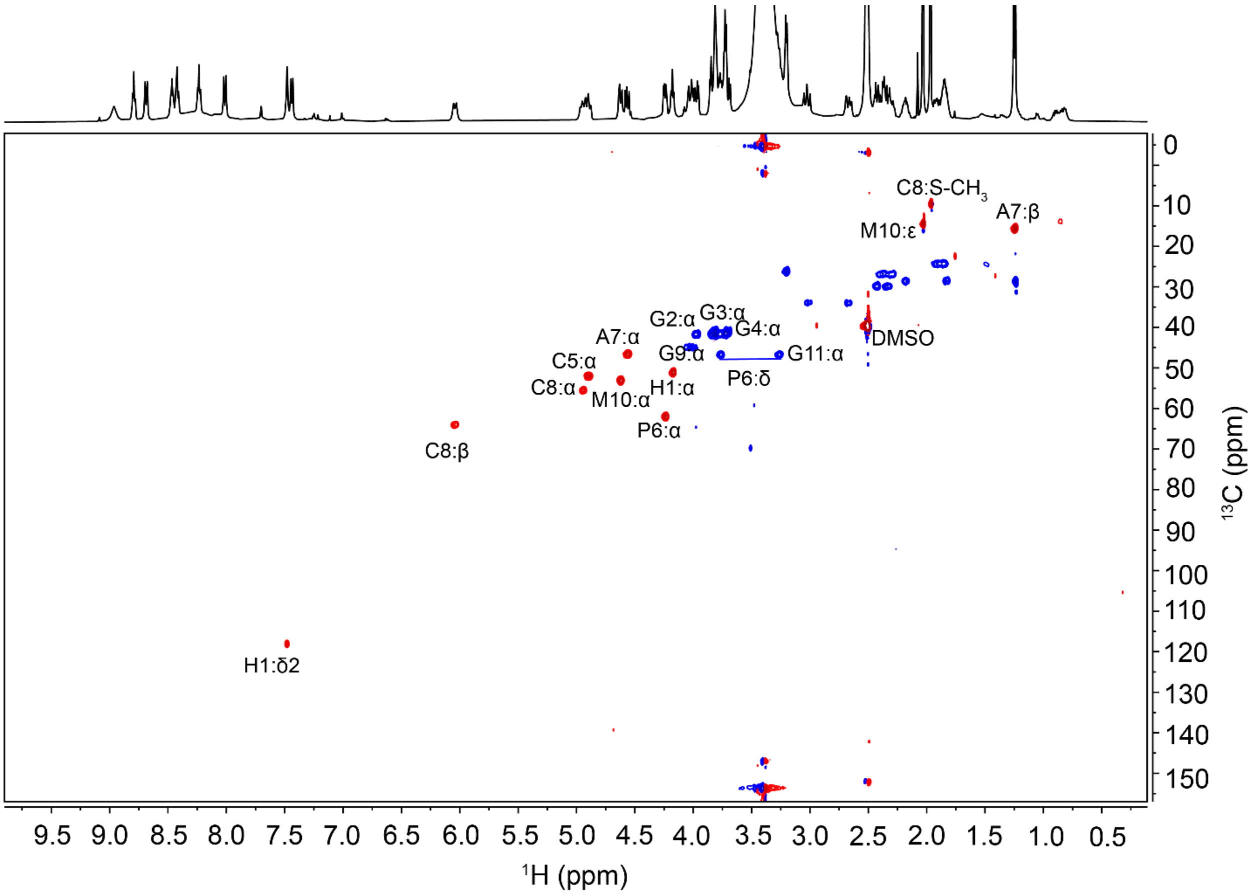
C8:S-CH3
A7:β
M10:ε
G2:α
G3:α
G4:α
A7:α
DMSO
G9:α
G11:α
C5:α
P6:δ
C8:α
M10:α
H1:α
P6:α
C8:β
H1:δ2
9.5 9.0 8.5 8.0 7.5 7.0 6.5 6.0 5.5 5.0 4.5 4.0 3.5 3.0 2.5 2.0 1.5 1.0 0.5
¹H (ppm)
0 10 20 30 40 50 60 70 80 90 100 110 120 130 140 150
¹³C (ppm)

**F**

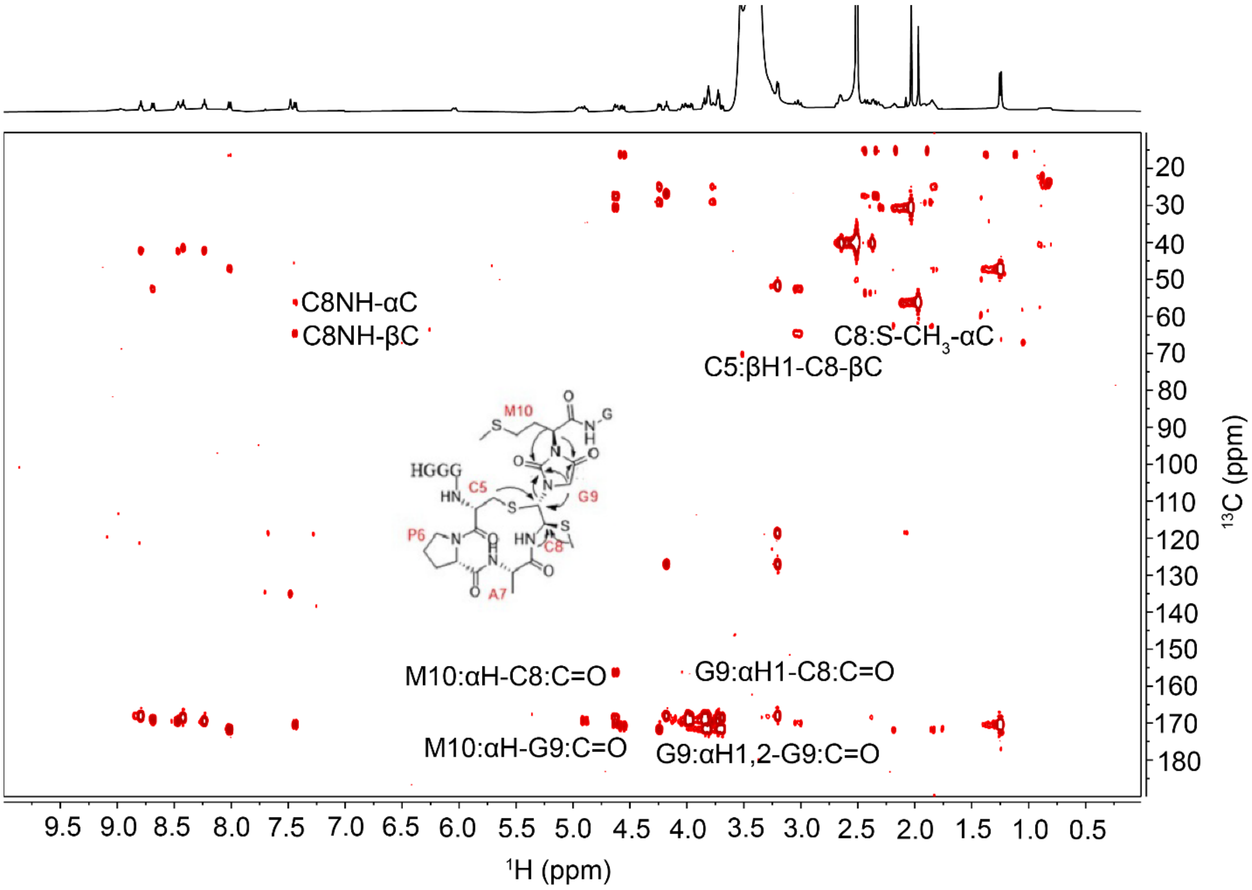
C8NH-αC
C8NH-βC
C8:S-CH3-αC
C5:βH1-C8-βC
M10
G
HGGG
C5
G9
P6
C8
A7
M10:αH-C8:C=O
G9:αH1-C8:C=O
M10:αH-G9:C=O
G9:αH1,2-G9:C=O
9.5 9.0 8.5 8.0 7.5 7.0 6.5 6.0 5.5 5.0 4.5 4.0 3.5 3.0 2.5 2.0 1.5 1.0 0.5
1H (ppm)
20 30 40 50 60 70 80 90 100 110 120 130 140 150 160 170 180
13C (ppm)

**G**

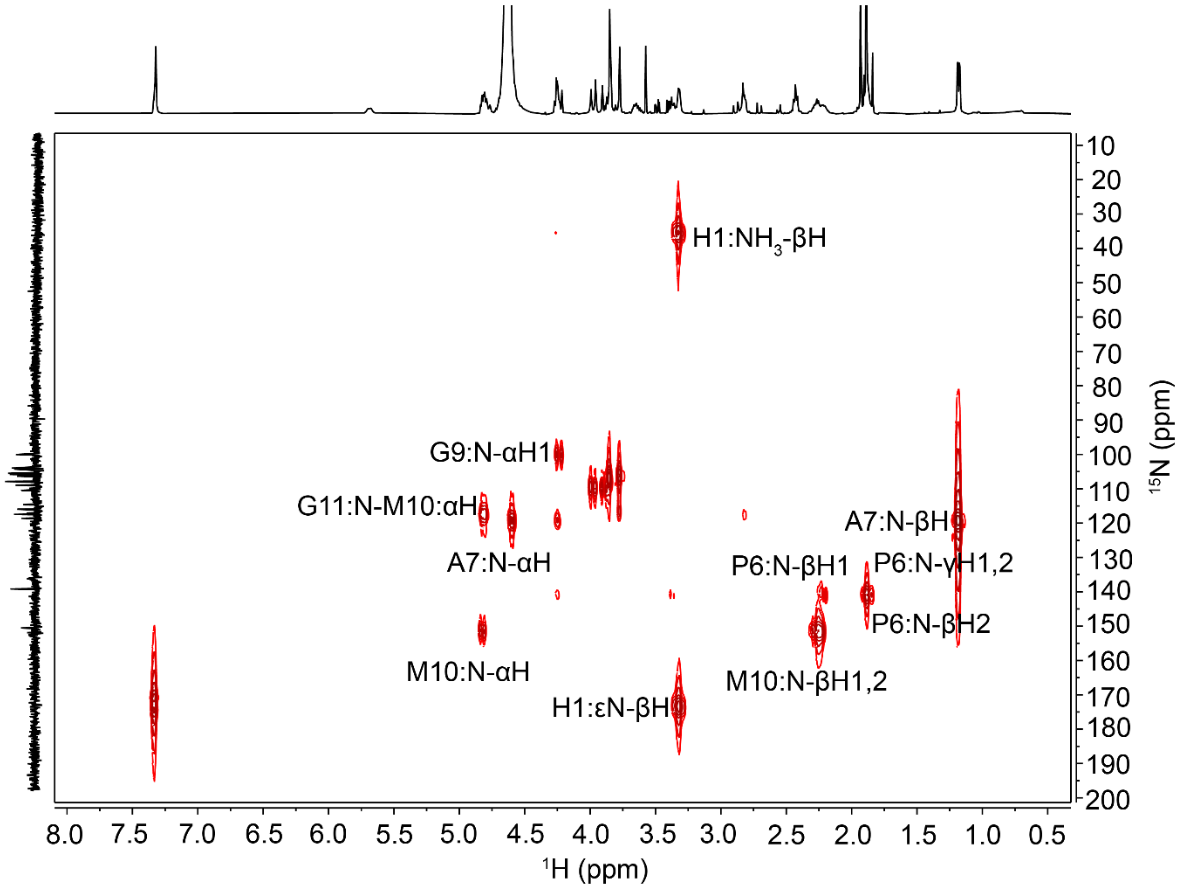

**Figure S6.** NMR spectra of LysC-cleaved ChrA* peptide in DMSO-d6 at 23 °C. Shown are: (A) $^{1}$H, (B) COSY, (C) TOCSY, (D) NOESY, (E) $^{1}$H-$^{13}$C HSQC, (F) $^{1}$H-$^{13}$C-HMBC, and (G) $^{1}$H-$^{15}$N-HMBC spectra.

**A**

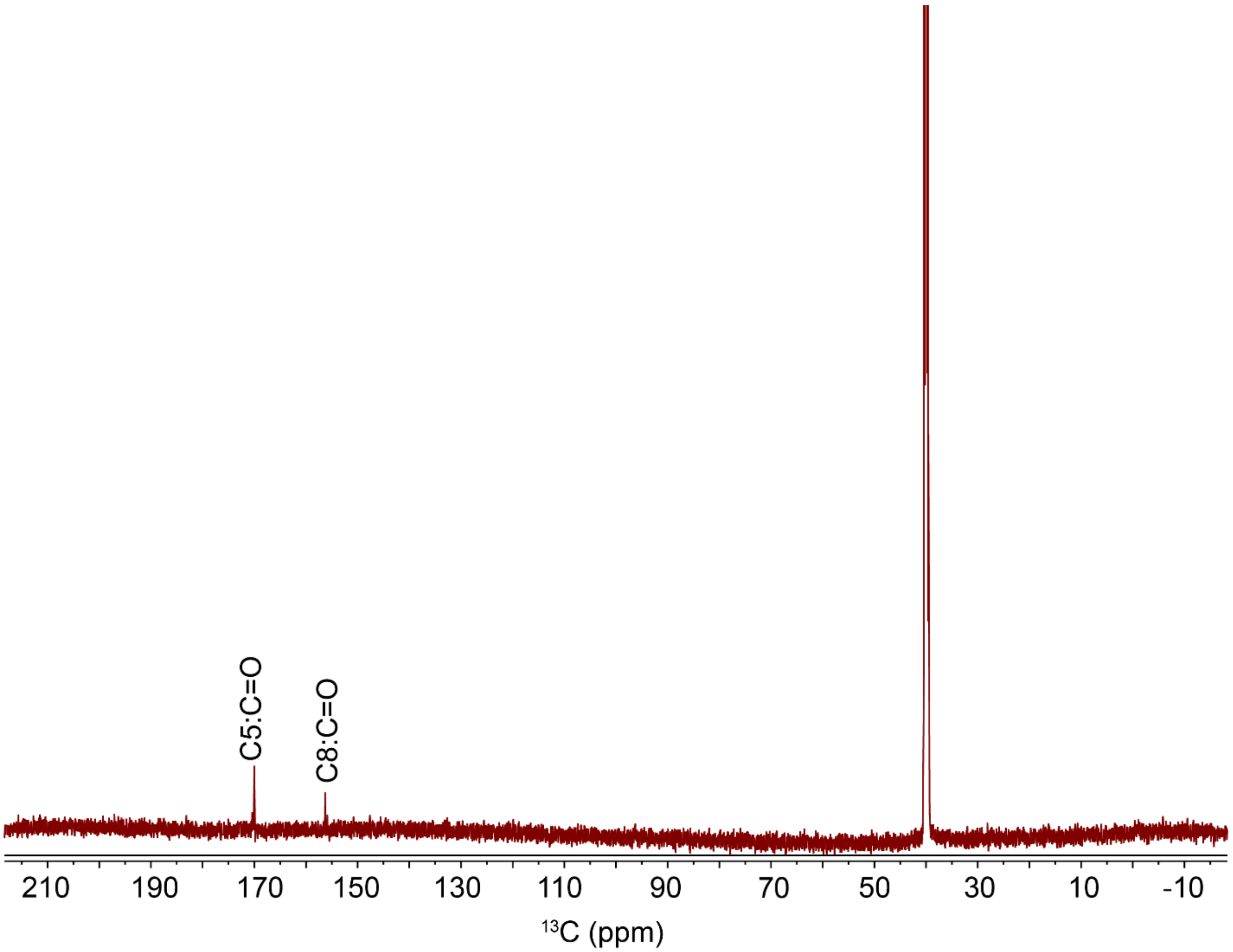

**B**

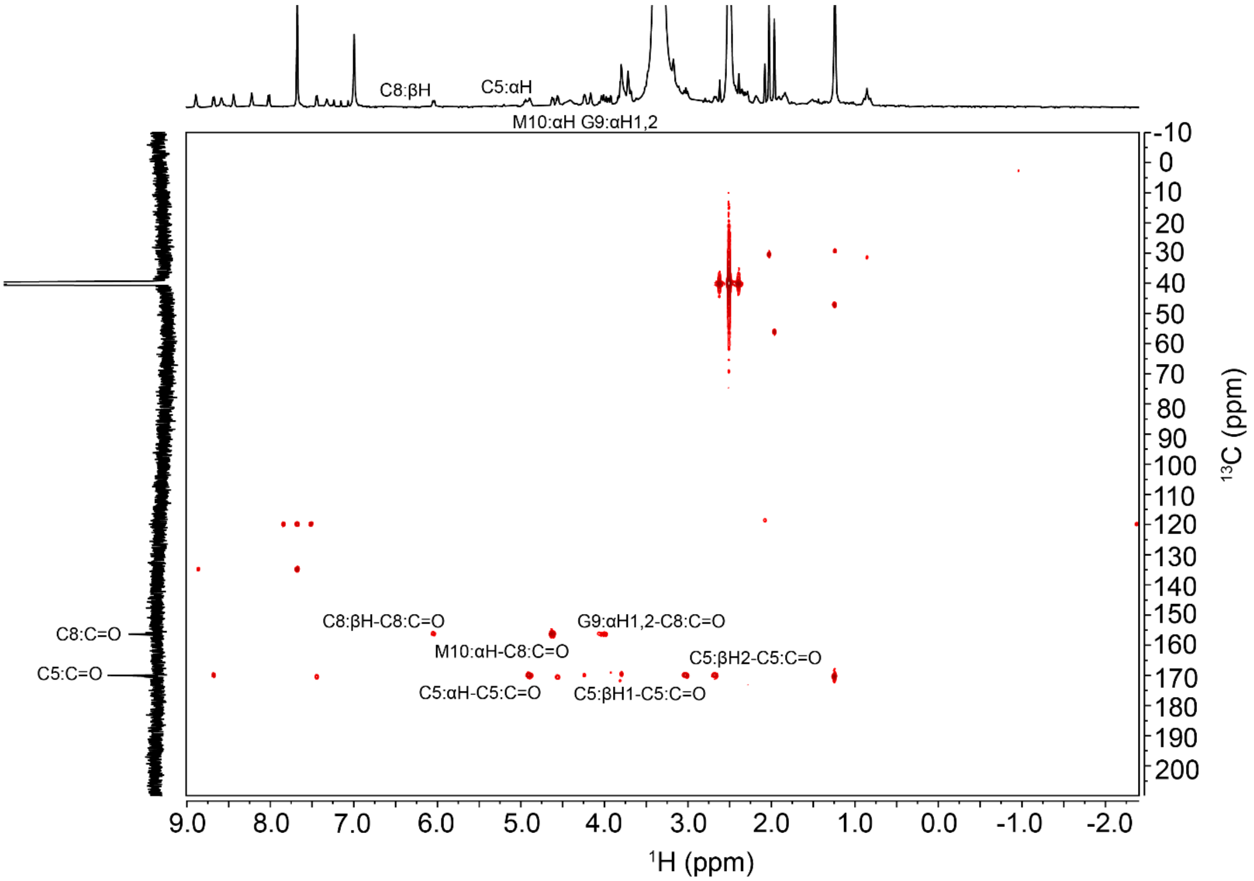

**Figure S7.** NMR spectra of LysC-cleaved 1-$^{13}$C-Cys-labeled ChrA* peptide in DMSO-d6 at 23 $^{o}$C. Shown are: (A) $^{13}$C and (B) $^{1}$H-$^{13}$C HMBC spectra.

**A**

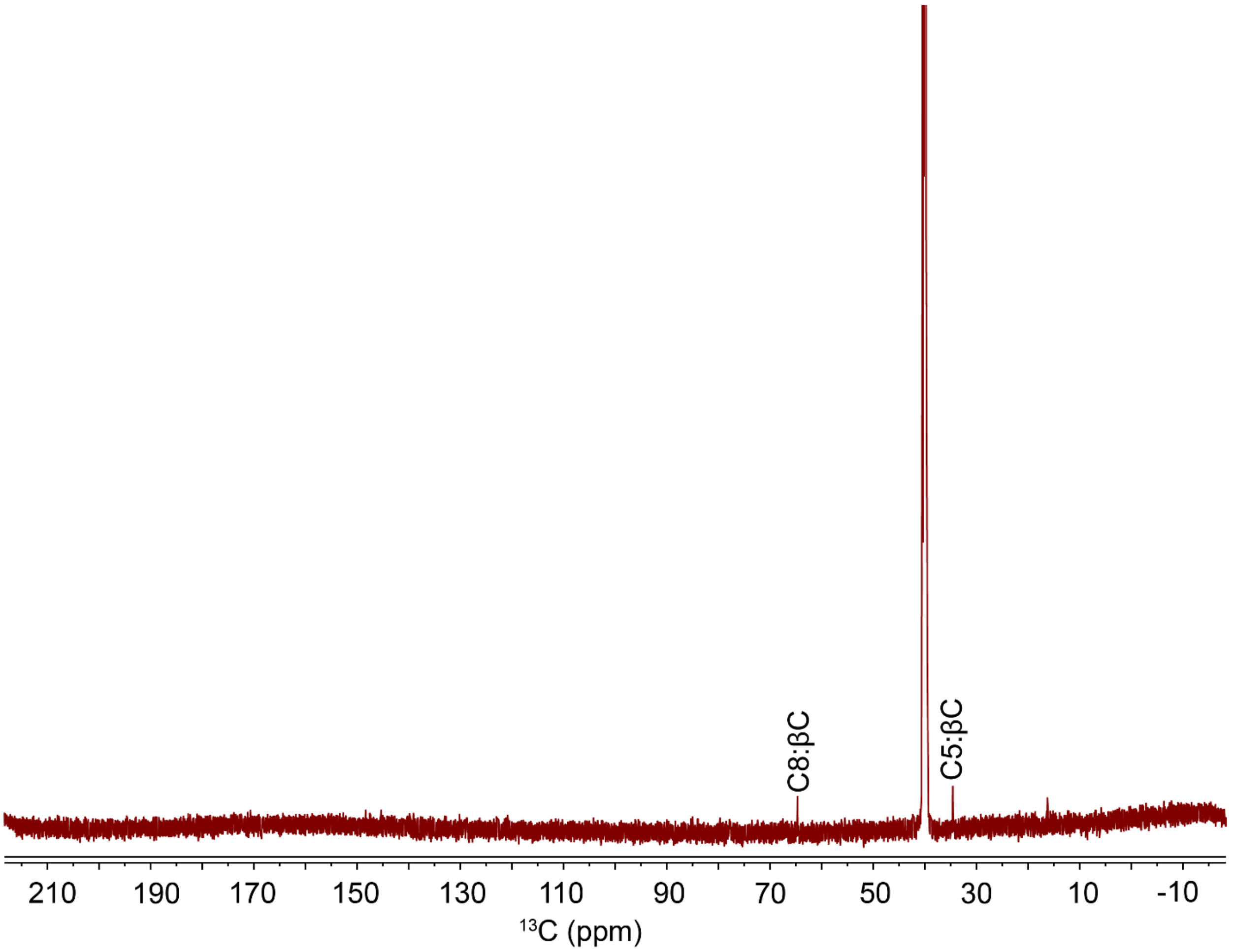

**B**

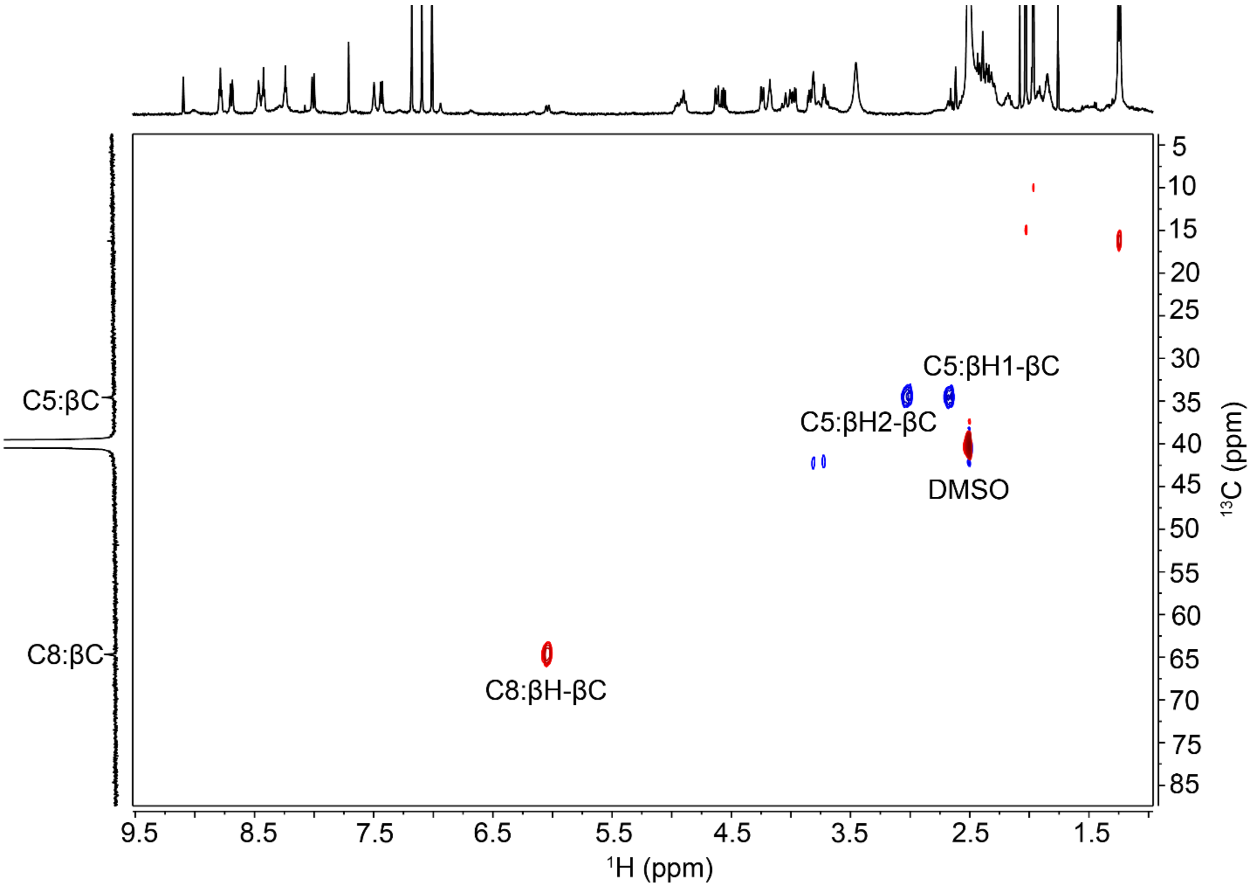

C5:βH1-βC
C5:βH2-βC
DMSO
C8:βH-βC
C5:βC
C8:βC
5
10
15
20
25
30
35
40
45
50
55
60
65
70
75
85
$^{13}$C (ppm)
9.5
8.5
7.5
6.5
5.5
4.5
3.5
2.5
1.5
$^{1}$H (ppm)

C

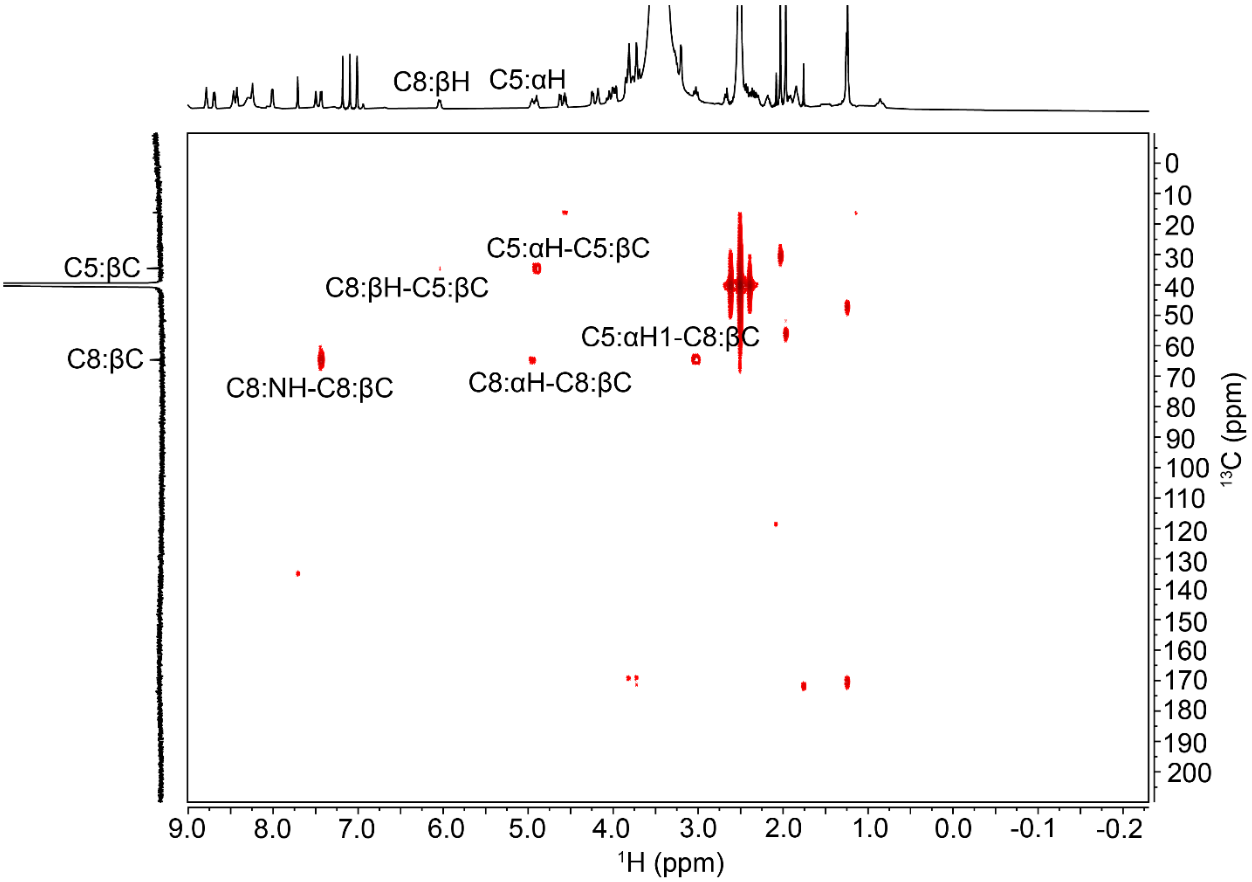

**Figure S8.** NMR spectra of LysC-cleaved 3-$^{13}$C-Cys-labeled ChrA* peptide in DMSO-d6 at 23 $^{o}$C. Shown are: (A) $^{13}$C, (B) $^{1}$H-$^{13}$C HSQC, and (C) $^{1}$H-$^{13}$C HMBC spectra.

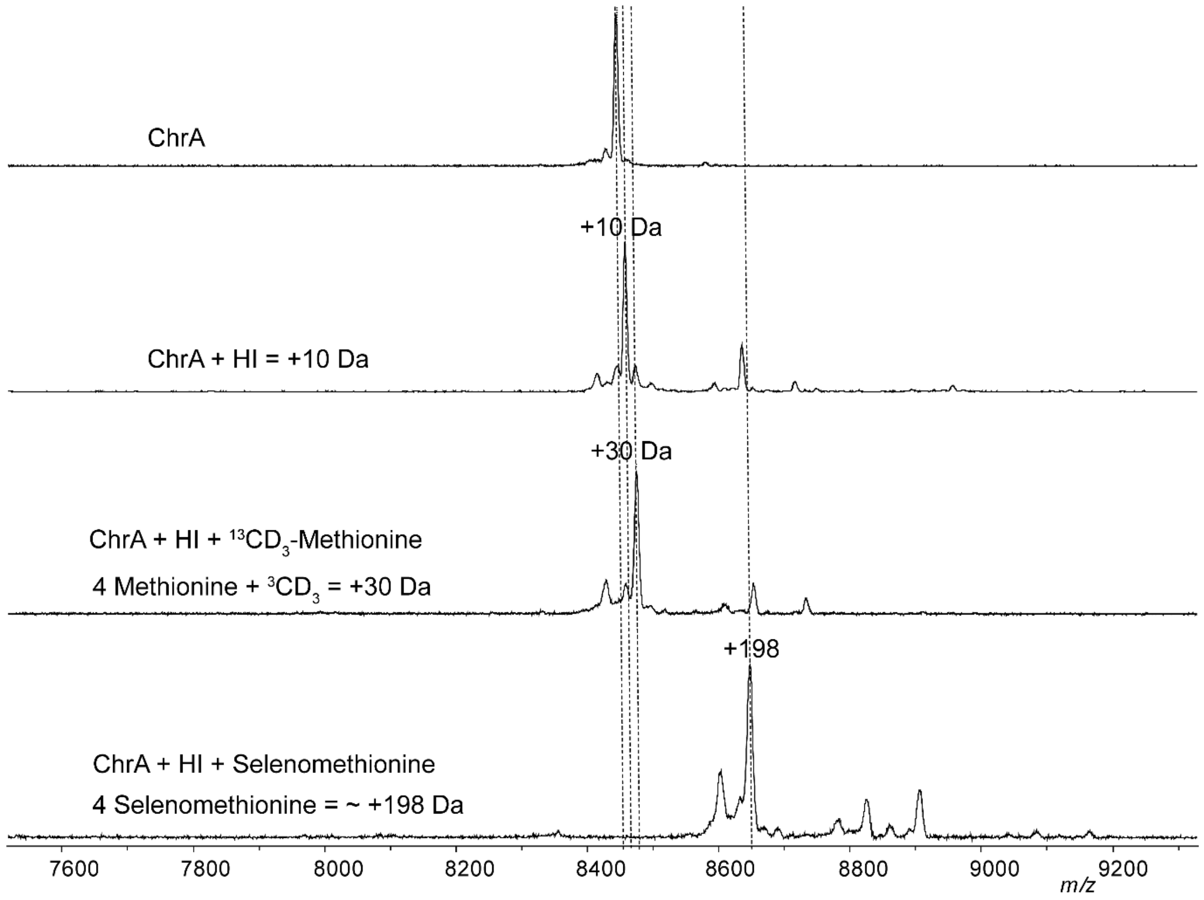

**Figure S9.** Determination of the methyl source for ChrHI catalysis. Shown are the MALDI-TOF mass spectra of ChrA produced by heterologous coexpression with ChrHI in M9 media supplemented with $^{13}CD_3$-methionine and selenomethionine. ChrA contains four Met residues (Figure 2A) and after ChrHI-modification one additional methyl group, resulting in a mass shift of 5 x 4 Da for each $^{13}CD_3$ group = 20 Da.

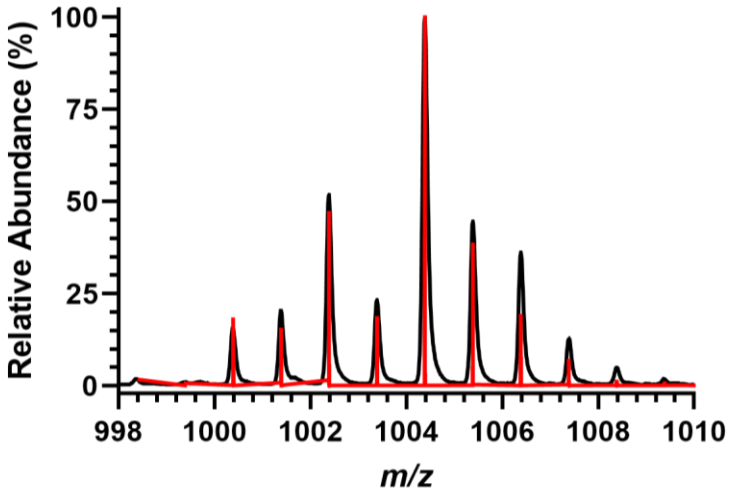

**Figure S10.** MALDI-TOF-MS analysis of selenoMet-ChrA*. The observed $[M+H]^+$ spectrum of the selenoMet-containing fragment (black) is overlaid on a simulated spectrum calculated by enviPat[9] (red). The isotope distribution confirms the presence of one selenium atom in the peptide.

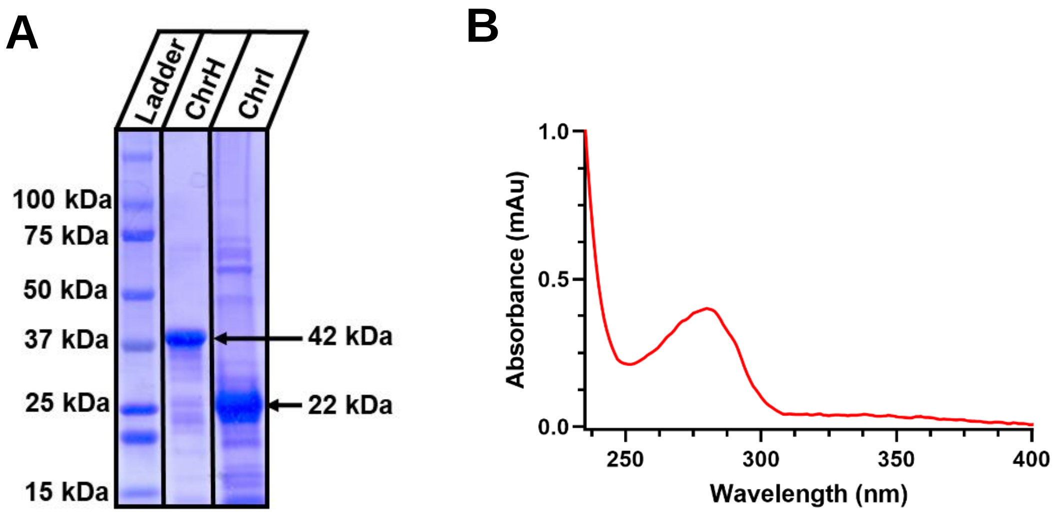

**Figure S11.** Preparation of ChrH and ChrI. ChrH and ChrI were purified and analyzed by SDS-PAGE. Lane 1 contains the Bio-Rad Kaleidoscope Prestained Protein Ladder standard, lane 2 contains $His_6$-ChrH, and lane 3 contains $His_6$-ChrI. (B) UV−vis spectral analysis of aerobically prepared ChrH.

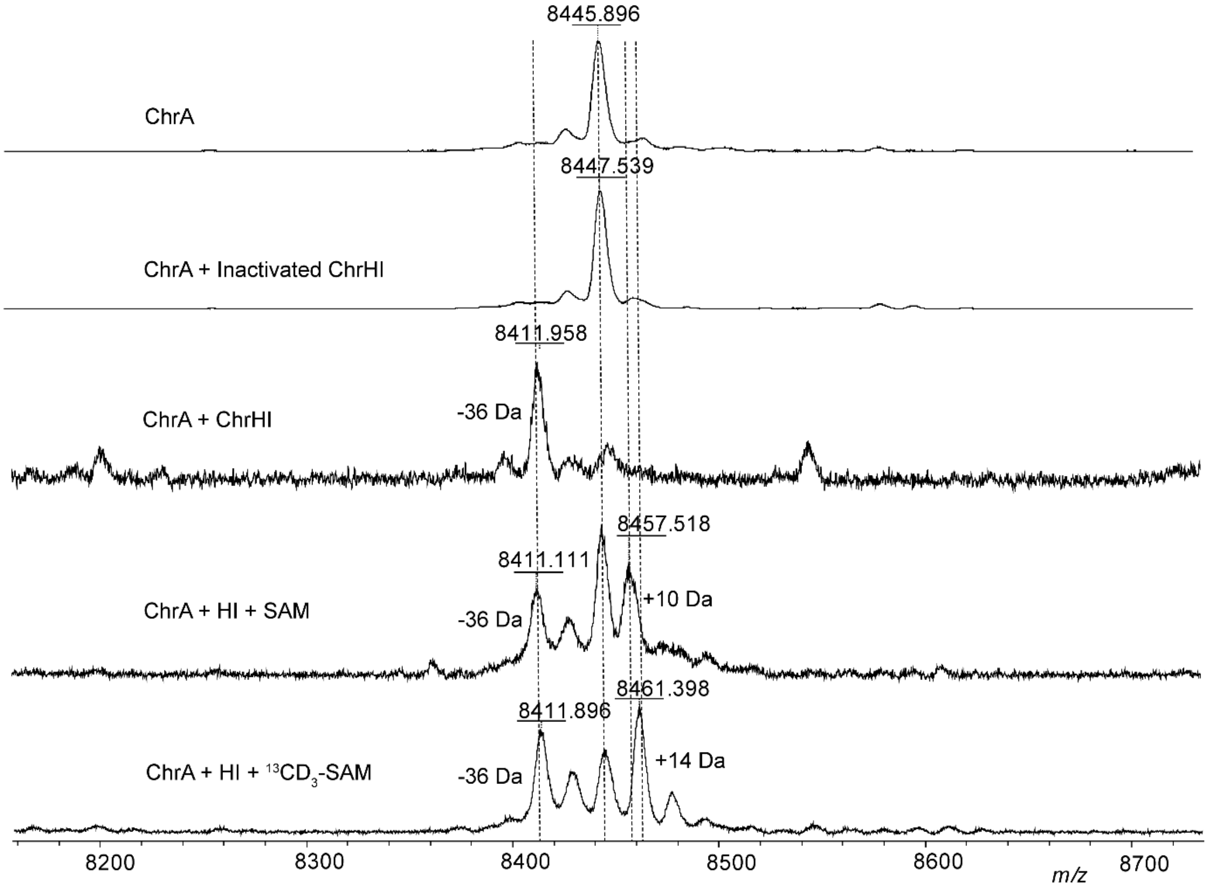

**Figure S12.** In vitro reconstitution of ChrHI. Shown are the MALDI-TOF mass spectra of ChrA, ChrA + denatured ChrHI, ChrHI reaction with and without SAM. The dashed lines indicate the expected *m/z* of modified and unmodified peptides.

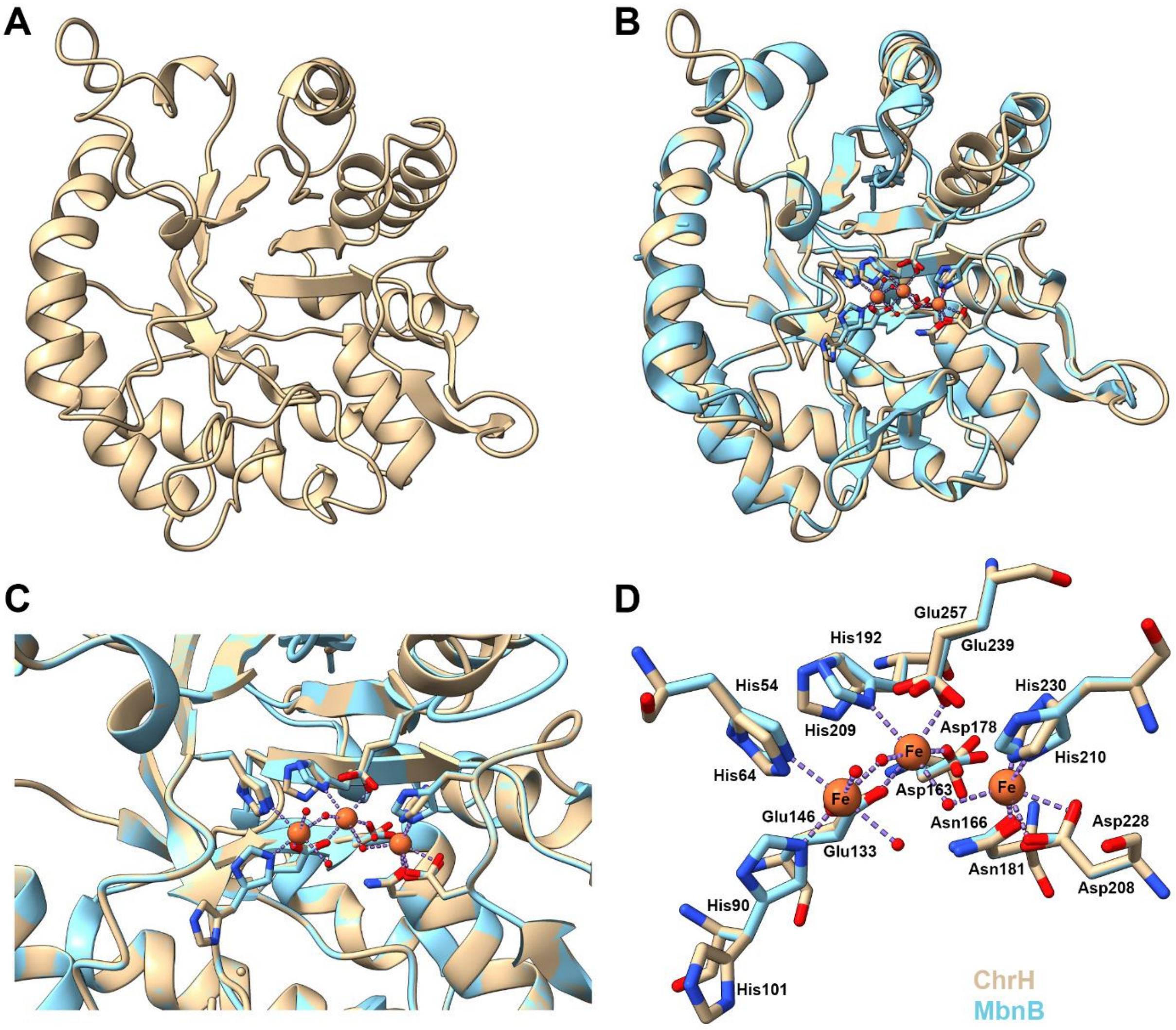

**Figure S13**. Predicted structure of ChrH using AlphaFold.[10] Shown are (A) the predicted structure of ChrH. (B) Predicted structure of ChrH (beige) superimposed with MbnB (aqua; PDB ID: 7TCX), (C) a zoom-in of the active site with Fe ions shown in MbnB, and (D) conserved residues that are likely to coordinate Fe ions.

A

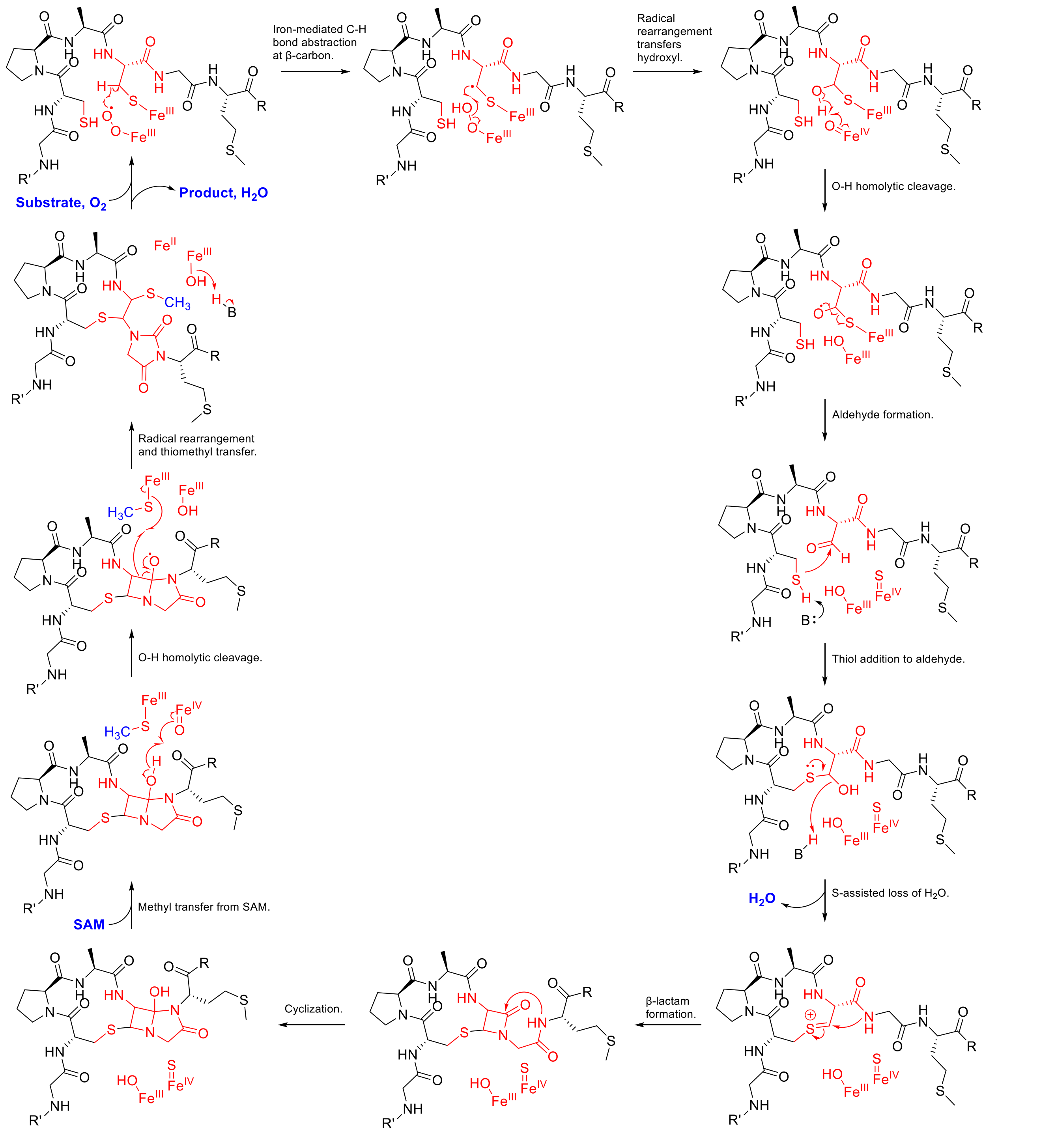

Iron-mediated C-H bond abstraction at β-carbon.
Radical rearrangement transfers hydroxyl.
O-H homolytic cleavage.
Aldehyde formation.
Thiol addition to aldehyde.
S-assisted loss of H2O.
H2O
β-lactam formation.
Cyclization.
Methyl transfer from SAM.
SAM
O-H homolytic cleavage.
Radical rearrangement and thiomethyl transfer.
Substrate, O2
Product, H2O

**B**

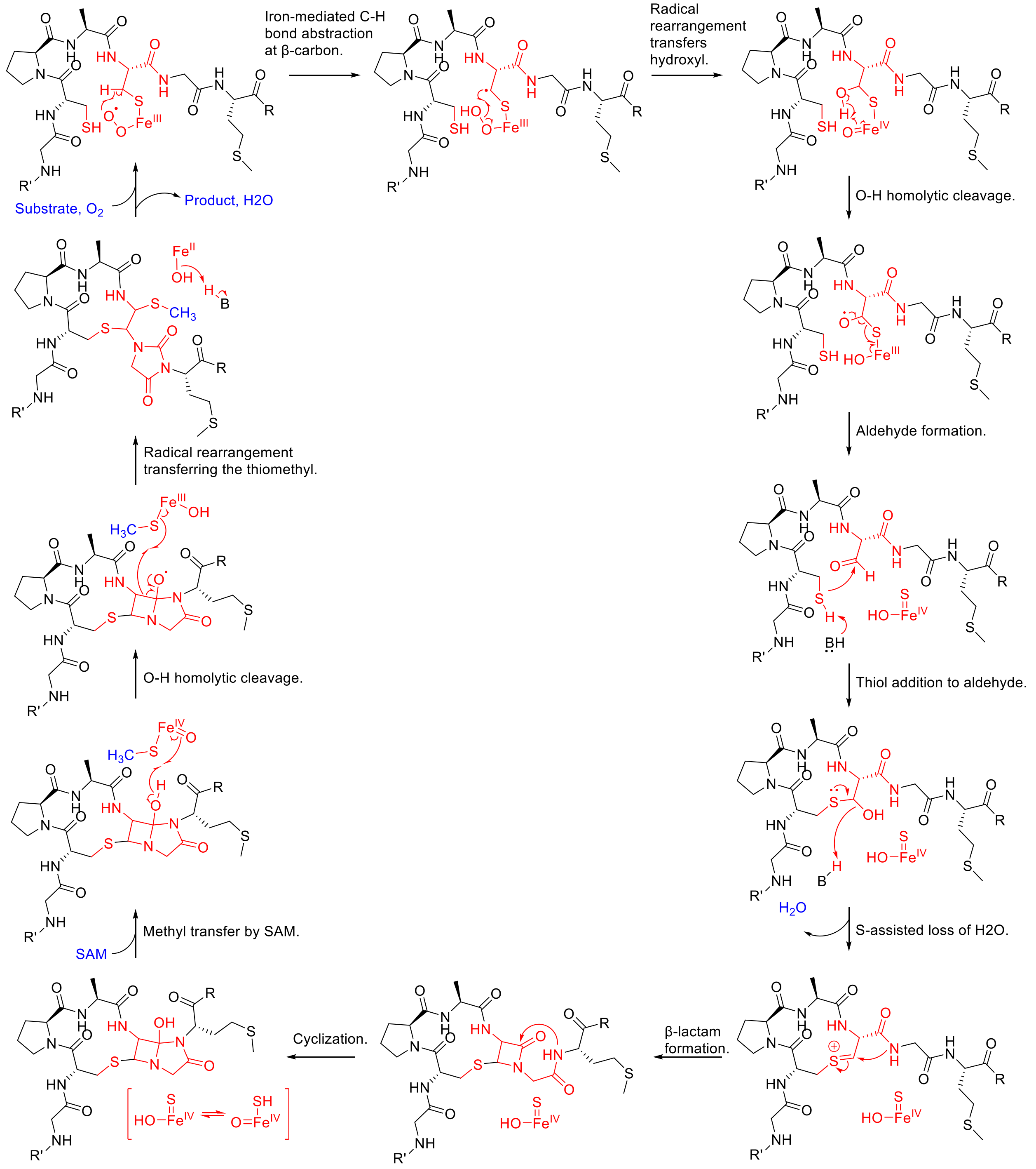
Iron-mediated C-H bond abstraction at β-carbon.
Radical rearrangement transfers hydroxyl.
O-H homolytic cleavage.
Aldehyde formation.
Thiol addition to aldehyde.
S-assisted loss of H2O.
β-lactam formation.
Cyclization.
Methyl transfer by SAM.
SAM
O-H homolytic cleavage.
Radical rearrangement transferring the thiomethyl.
Substrate, O2
Product, H2O

**C**

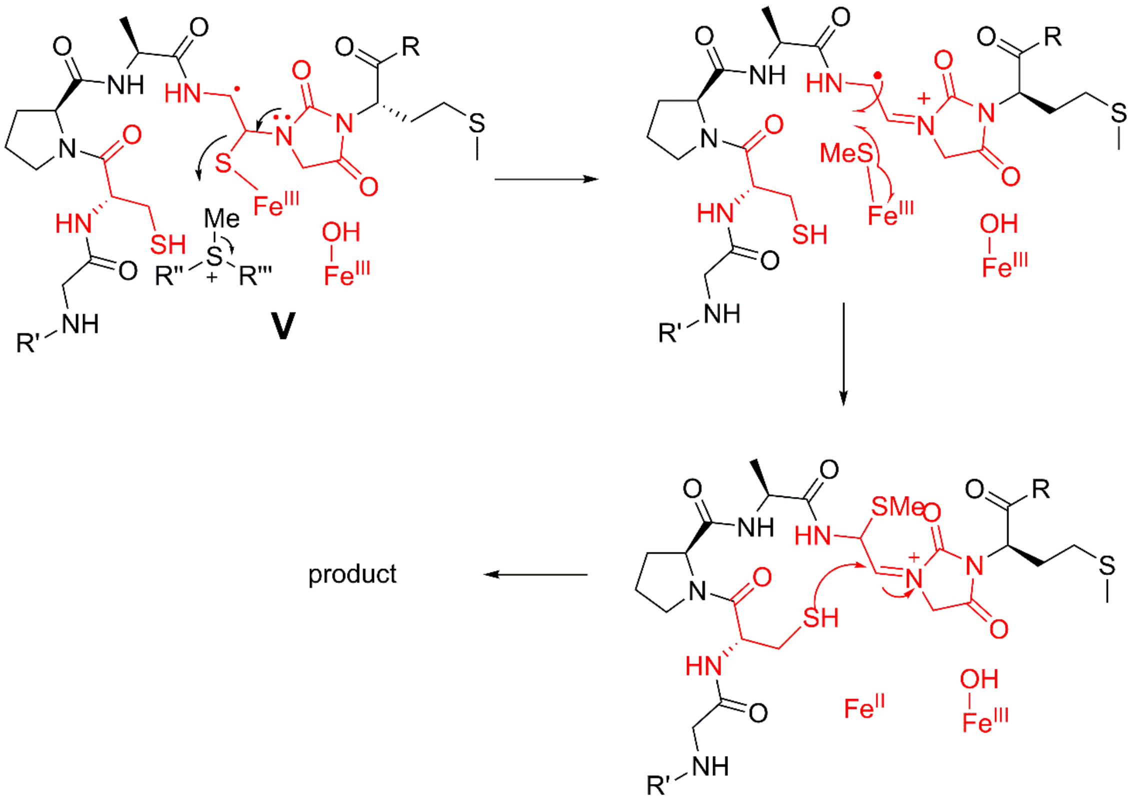

**Figure S14.** Alternative mechanisms for ChrHI catalysis. (A) Mechanism involving mixed valent diiron form of the enzyme in which the Cβ-S bond is broken early in the pathway and the macrocycle is formed prior to the imidazolidinedione formation. The mechanism also features several other alternative steps (compared to Figure 6) that may form the new bonds in the product. (B) Mechanism in which a single iron is involved in the chemistry (a second or third iron could be involved in stabilizing charged intermediates or positioning substrate and/or intermediates). (C) Mechanism in which the carbon-sulfur bond involving the thiomethyl group is formed by radical rather than heterolytic chemistry. This mechanism has some resemblance to thiomethylation by radical SAM proteins in that an iron-bound sulfur atom is methylated by SAM and then transferred to an intermediate radical.[11-16] Many hybrid mechanisms can be drawn that take aspects of the mechanisms in Figure 6 and S14A-C. Given that currently very little is known about intermediates and substrate positioning, these alternative hybrid mechanisms are not drawn here.

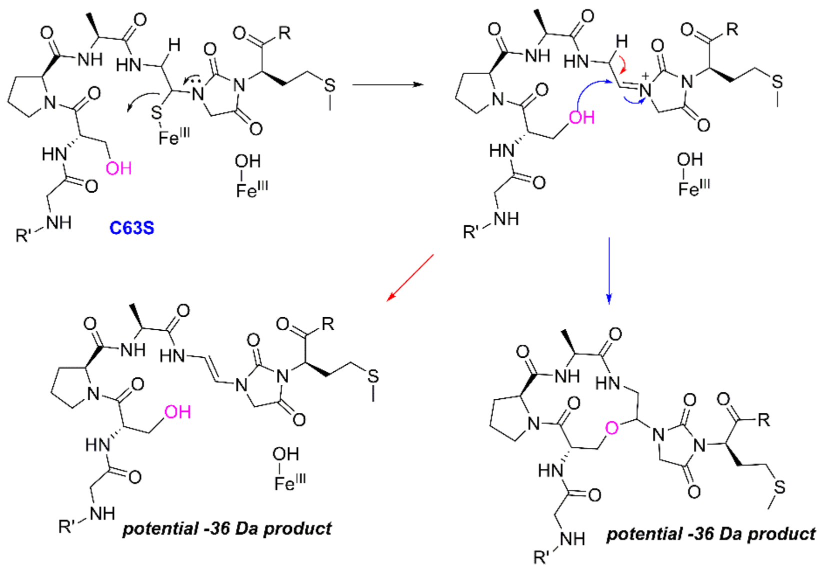

**Figure S15.** Possible explanation for the formation of a -36 Da product from the ChrA-C63S variant. This mechanism is based on that shown in Figure 6 of the main text for the WT reaction. The mechanisms in Figure S14 can be similarly used to explain formation of a -36 Da product with this variant.

**Table S1.** Chemical Shifts of ChrA in 90% $H_2O$ and 10% $D_2O$ at 25 $^oC$. $^1H$ chemical shifts were referenced to water peak at 4.7 ppm, and $^{13}C$ was referenced to TMS at 0 ppm.

| number | AA | NH | αH/αC | βH/βC | γH/Cγ | δH | others |
|---|---|---|---|---|---|---|---|
| 1 | H | | 4.26<br>54.4 | 3.31,28.0 | HD2:7.32,121.0<br>HE1: 8.53,136.8 | | |
| 2 | G | 8.71 | 3.95<br>44.3 | | | | |
| 3 | G | 8.36 | 3.90<br>44.6 | | | | |
| 4 | G | 8.29 | 3.83<br>44.8 | | | | |
| 5 | C | 8.24 | 4.81<br>53.1 | 3.22<br>39.7 | | | |
| 6 | P(trans) | | 4.23<br>65.0 | 2.24, 1.84<br>31.0 | 1.99,1.88<br>26.8 | 3.65,3.57<br>50.0 | |
| 7 | A | 8.177 | 4.378<br>51.5 | 1.274 | | | |
| 8 | C | 7.846 | 4.44<br>60.3 | 3.36, 2.96<br>36.5 | | | |
| 9 | G | 8.264 | 3.84<br>44.8 | | | | |
| 10 | M | 8.067 | 4.41<br>54.9 | 2.02, 1.88<br>32.4 | 2.50, 2.41<br>31.4 | 1.974<br>16.1 | |
| 11 | G | 8.20 | 3.72<br>44.7 | | | | |

**Table S2.** Chemical Shifts of ChrA* in DMSO-d6 at 23 $^oC$. $^1H$ and $^{13}C$ chemical shifts were referenced to 2.5 ppm and 40 ppm for DMSO, respectively. $^{15}N$ chemical shifts were referenced externally to liquid ammonia at 0 ppm.

| number | AA | NH/$^{13}C$=O | $^{15}N$ | αH/$^{13}C_\alpha$ | βH/$^{13}C_\beta$ | γH/$^{13}C$ | δH<br>others |
|---|---|---|---|---|---|---|---|
| 1 | H | 168.1 | 39.2<br>($NH_3$) | 4.17<br>51.8 | 3.19<br>26.7 | HD2:7.48, 118.6<br>HE1: 8.97, 134.6,<br>C[illegible]:127.2 | |
| 2 | G | 8.78<br>169.6 | 109.5 | 3.98, 3.82<br>42.3 | | | |
| 3 | G | 8.46<br>169.5 | 107.4 | 3.81<br>42.5 | | | |
| 4 | G | 8.23<br>169.1 | 105.3 | 3.72<br>42.4 | | | |
| 5 | C | 8.68<br>170.0 | 116.8 | 4.89<br>52.5 | 3.02.2.67<br>34.4 | | |
| 6 | P<br>(trans) | 171.8 | 139.2 | 4.23<br>62.6 | 2.17, 1.83<br>29.0 | 1.91, 1.85<br>24.9 | 3.77,3.26<br>47.2 |
| 7 | A | 8.01 | 115.4 | 4.56 | 1.24 | | |

| | | | | | | | |
|---|---|---|---|---|---|---|---|
| | | 170.5 | | 47.1 | 16.2 | | |
| 8 | C | 7.44<br>156.5 | 118.0 | 4.94<br>56.1 | 6.04<br>64.6 | (-S-$CH_3$)<br>1.96,<br>10.0 | |
| 9 | G | Losing NH<br>170.3 | 100.4 | 4.04, 4.01<br>45.5 | | | |
| 10 | M | Losing NH<br>168.7 | 150.6 | 4.62<br>53.6 | 2.37, 2.30<br>27.4 | 2.43,2.34<br>30.4 | 2.03,<br>15.0 |
| 11 | G | 8.41<br>171.6 | 105.1 | 3.81, 3.71<br>41.5 | | | |

**Table S3:** Predicted internal ChrA* fragments and the corresponding bonds cleaved.

| | |
|---|---|
| 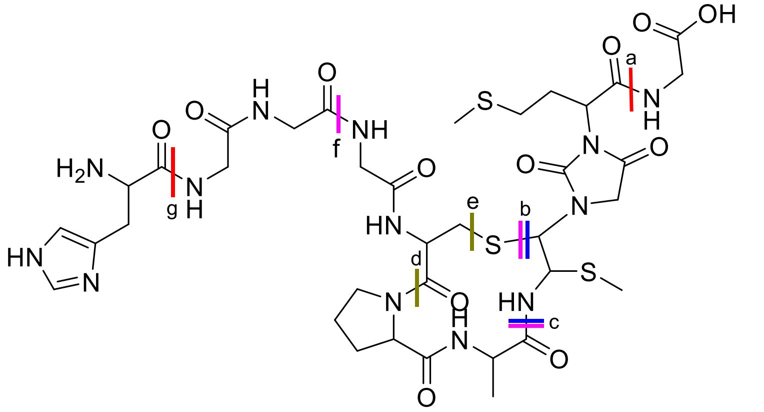<br>Parent structure | |
| **Internal fragment** | **Bonds cleaved** |
| 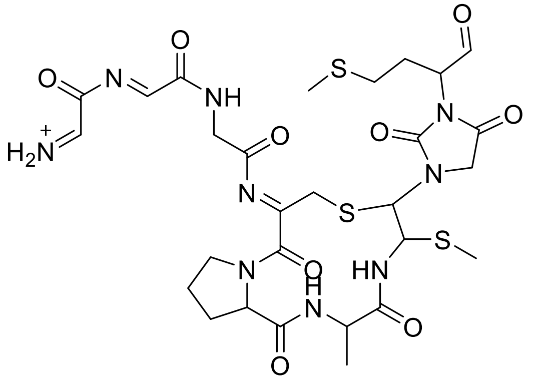<br>calc'd m/z: 740.1950, observed m/z: 740.1966 | a and g |

| | |
|---|---|
| 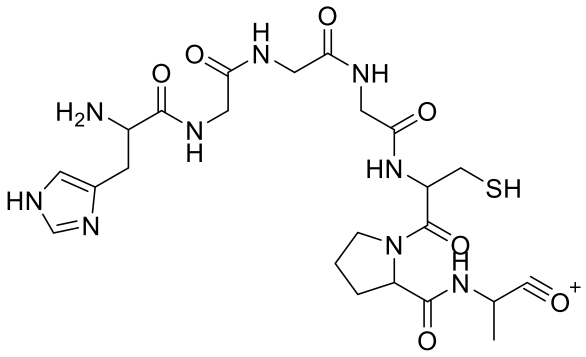 calc'd m/z: 580.2297, observed m/z: 580.2309 | c and b |
| 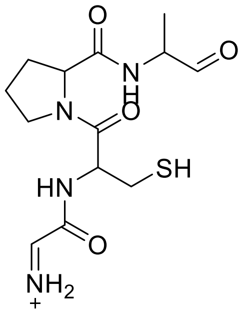 calc'd m/z: 329.1279, observed m/z: 329.1269 | b, c and f |
| 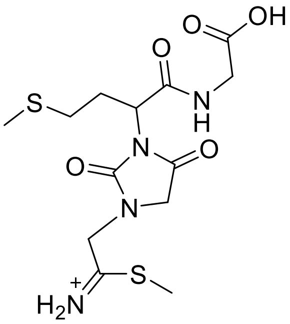 calc'd m/z: 377.0948, observed m/z: 377.0957 | b and c |
| 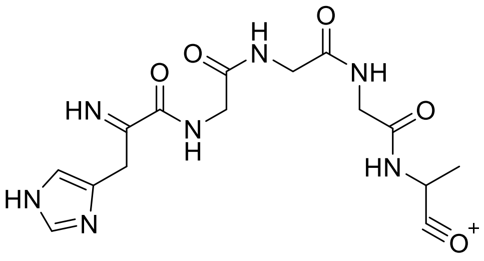 calc'd m/z: 378.1521, observed m/z: 378.1530 | d and e |

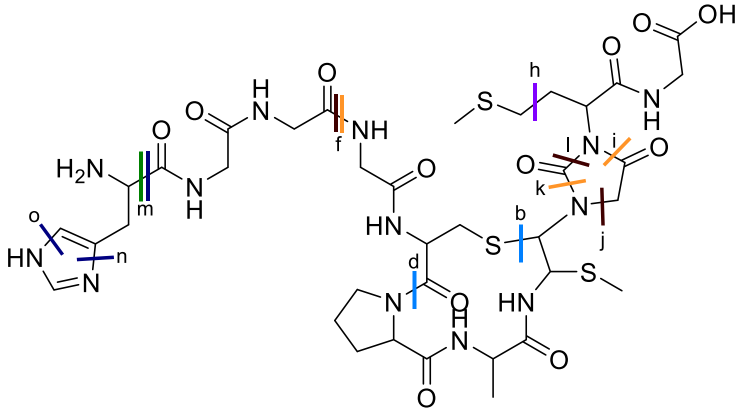

Parent structure

| Internal fragment | Bonds cleaved |
|---|---|
| 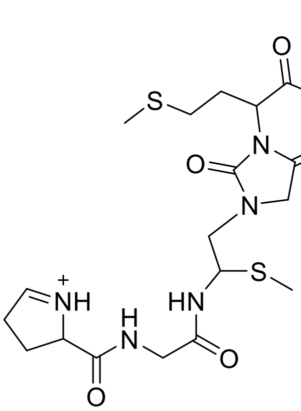 calc'd m/z: 531.1691, observed m/z: 531.1702 | b and d |
| 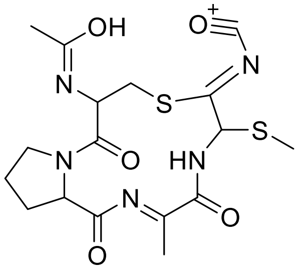 calc'd m/z: 440.1057, observed m/z: 440.1046 | f, i and j. Deamination of the N-terminal amine. |
| 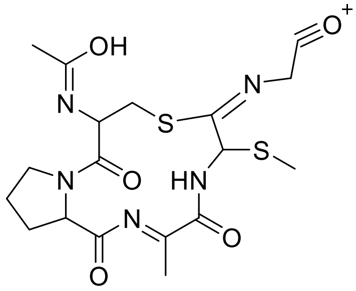 calc'd m/z: 454.1214, observed m/z: 454.1224 | k, f and i. Deamination of the N-terminal amine |

| | |
|---|---|
| 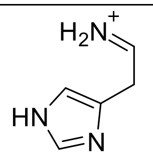<br>calc'd m/z: 110.0713, observed m/z: 110.0716 | m |
| 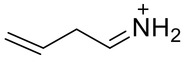<br>calc'd m/z: 70.0652, observed m/z: 70.0656 | n, m and o |
| 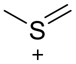<br>calc'd m/z: 61.0107, observed m/z: 61.0112 | h[17] |

**Table S4.** Sequences of *E. coli* codon-optimized genes of the *chr* BGC used in this study.

| **Gene Name** | **Sequence (5'à3')** |
|---|---|
| *chrA* | ATGAAGATTCCGGCTCTGGTGATGGCCAGCCTGCTGGCGGTATCAGTGAGCGG<br>GCAAACCACGAAACCGGTCAAAAAAGGCACGAAAAGTGTTAAAAAAGTGAAA<br>AAAATGGATAGCCCAAAAACCGTTAAAGCAGAACCCACCAAAGTCGTTAAAC<br>GTGACACGATCCTGAAACATGGCGGCGGTTGTCCGGCGTGCGGTATGGGGTGA |
| *chrH* | ATGAAGAAACCTCTCCTTGGATTAGCCATGATGCCGGAAGCTGACTTTGTTTCC<br>GCTATTCTTCCGCTTTTGCAAACGCAAAGTGTTGACGTACTCGAGTGGAGCTTC<br>GATACGCTGTATGACGTCGAAGAGCCGGAATGGCTGTCCGGACTGCTTGACTT<br>CTATTCAGACAATTCACGCCTGTTGGGACACGGGGTGTACTATTCATTATTCGA<br>CGCCCGCTGGATGGAGCGCCAAGAGATTTGGCTCAAGAAGCTTAAAGAAGAG<br>GTGAAGCGCCGCAAGTACAATCATATCACCGAGCATTTTGGCTTCATGAACAC<br>TGAGAATTTCCACCAAGGCGTACCTCTTCCTGTACCACTTCTGCCGAAGACGCT<br>GCAAATTGGGAAGGATCGCCTGTCTCGTCTCCAGGACGCCGTGGAGATCCCGG<br>TGGGTGTCGAGAATCTGGCATTCTCTTTCAGCATGGATGACGTCAAGGAGCAA<br>GGTGAATTCTTGGACCGTTTGGTGGAAGACATCGACGGGTTCCTCATCCTCGAT<br>CTCCACAATATTTACTGCCAGAGCTGCAACTTCAAGGTGGACATGATGGAGAT<br>CATCAATTTATATCCGCTCGAGAAGGTCAACGAGATCCACCTCTCCGGCGGCT<br>CTTGGCAAGAGTCGGCATACGGCAAGAAACCTGTGCGCCGTGATACGCATGAC<br>GACCGTATCCCTGAAGAGATTCTGAATTTATTACCCGAGGTTTTAATTCAGTGC<br>CATCCTGAGTACGTAATTATTGAACGCTTAGGCCACACACTTAATACAGAGGT<br>GGCCAAGCAAATTTTCTTCGATGACTTTAACCGCGTAAAGAAAATCTTGGAGT<br>TATCAGATTACCCAGTCGGAGAAGAGAAAATTTGGAATCGCCAGAATACCGTC<br>CACTCGAAGCCTGTTGAAGATCTCCTCTTGTATGCTGAGCAAACTGAATTAACC<br>CGTTTACTTTTCGATGGTAATACGACGGAATCTATTAAGAATCGCGATTTTCAT<br>TATTTCAAGCCTGAATCCTGGGACGAGGAGATGATCACAACTGCGCAACAGAT<br>CATTAAGAAGTGGAACCCGTAA |
| *chrI* | ATGAAAATGACTACTGTATTTCTGCTGATTGATTACCGCGGTTTTAACAGCATT<br>GATCGCTGGGCTGTTCTACGCTTACTCATGCTCAGTGGTACTTGGCTTAGGCAA<br>GTTAAGTGACACGGAGTATCTTAAAAGCATGCAAAATATCAATCGTGAAATCT<br>TGAACCCTGTTTTCTTCATGAGCTTCATGGGTACCGCCGTTCTCTTACCAGTTGC<br>GACGTTTCTTTTCCGTGGTGAGCAACCTTCGTTCTTATTCTTACTGTTAGCCTCG<br>GCCGCGTACCTGATTGGCGTGTTTGGTGTTACGATCGTAGGCAACGTCCCGTTA |

| | |
|---|---|
| | AATGATATGTTGGACAAATTTGATATCTCCGGTTCAACCATGGATGTAGTCAA GCAAATGCGCGACCGTTTCGAGAACCGCTGGAATCTCCTGAATAACGTGCGTA CTGTGTTTTCAGTAATTTCTATCGCATTAGTGGTTTGCGCGTGTATTTGGAACC GCCAGCTGACGTAA |

**Table S5.** Primers used for cloning of *chr* genes.

| Primer Name | Primer sequence (5’à3’) |
|---|---|
| ChrA-F | ATCACCATCATCACCACAGCCAGATGAAGATTCCGGCTCTGGTGATG |
| ChrA-R | CATTATGCGGCCGCAAGCTTCACCCCATACCGC |
| ChrH-F | ATAAGGAGATATACATGAAGAAACCTCTCCTTGGATTAGCCATGCTGCCGGAAG |
| ChrH-R | CAGGCGCGCCGTTACGGGTTCCACTTCTTAATGAT |
| ChrI-F | TGGCAGATCTCATGAAAATGACTACTGTATTTCTGCTGATTACCGCGG |
| ChrI-R | AGCAGCCTAGTTACGTCAGCTGGCGGTT |
